# View Tomo: Context-aware targeting and analysis in electron cryo-tomography

**DOI:** 10.64898/2026.04.21.719727

**Authors:** Robert Gebauer, Emily A. Machala, Yuliia Mironova, Märit-Runa Jönsson, Jędrzej Mazur, Clara A. Feldmann, Marcel A. Zimmeck, Emma Silvester, Enrico Caragliano, Julian Falckenhayn, Enoch Lok Him Yuen, Tarhan Ibrahim, Jan Hellert, Tolga O. Bozkurt, Rainer Kaufmann, Emmanuelle R. J. Quemin, Kay Grünewald, Vojtěch Pražák

## Abstract

Electron cryo-tomography (cryoET) resolves cellular structure in three dimensions, yet region selection is still typically based on two-dimensional projection images. Here, we introduce View Tomo, a workflow for rapid acquisition of low-magnification tomograms that enables screening, targeting and analysis in 3D. View Tomo tilt series are acquired in minutes at low dose (~3 e^−^/Å^2^), producing high-contrast tomograms that remain compatible with subsequent high-resolution structural determination. We implemented View Tomo using an automated acquisition and reconstruction pipeline for rapid alignment. Across multiple viral and cellular systems, view tomograms revealed membrane remodelling events, assembly intermediates and cellular organisation that are difficult to identify in projection images. These data enabled targeted high-resolution imaging and quantitative analysis of spatial relationships within cells. View Tomo therefore extends cryoET workflows by improving target selection, enabling analysis of mesoscale organisation, and facilitating integration with correlative imaging approaches.

## Main

Cryogenic electron microscopy (cryoEM) is a central approach in structural biology, enabling analysis of biological specimens across a wide range of scales and experimental contexts. The three most widely used modalities of cryoEM are: single particle analysis, micro-electron diffraction (microED) and electron cryo-tomography (cryoET). Single particle analysis and microED are primarily used for structural determination of purified macromolecules, whereas cryoET enables structural studies of heterogeneous specimens across scales, from molecular assemblies to cells. CryoET in particular has transformed our ability to examine the structural organisation of macromolecules within their native cellular environment^1^. By resolving the complex spatial relationships between proteins, membranes, and organelles in three dimensions (3D), it offers a window into the inner life of a cell at molecular resolution and under native conditions^2–5^. Combined with complementary techniques, it is central to integrative structural biology, where multiple data types are brought together to analyse macromolecular organisation, conformational heterogeneity and structure-function relationships *in situ*^*6*^.

In recent years, the range of specimens accessible to cryoET has been greatly expanded by new approaches such as focused ion beam (FIB) milling^7^ and advances in cryo correlative light and electron microscopy (cryoCLEM)^8^. With cryoFIB milling, thinning of vitrified specimens leads to electron-transparent sections called lamellae from regions that would otherwise be too thick under the transmission electron microscope (TEM)^9^. CryoCLEM enables localisation of specific biological structures using fluorescent signals, guiding not only lamella placement during FIB milling but also the subsequent selection of regions of interest (ROI) for high-resolution tomogram acquisition and the interpretation of data *a posteriori*. These capabilities are continuing to evolve, especially with the development of super-resolution cryoCLEM approaches, in which individual fluorescent proteins can in principle be localised with nanometre-scale precision^10–13^. However, due to the challenges of achieving high correlation accuracy by multi-modal imaging, reliable identification of specific structures for cryoET acquisition often still depends on the capacity to visualize them in low-magnification projection images.

Relying on projection images for both screening and targeting is a fundamental limitation of current cryoET practice, requiring not only experience but a degree of serendipity. Projection images can be sufficient to identify large structures such as mitochondria, but this relies on interpreting membrane densities and is therefore not always reliable. Because membrane contrast is strongly orientation-dependent, polar regions of such structures are often difficult to recognise. This is further compounded by the fact that projection images, even of very thin lamellae (~150 nm in thickness or less), contain multiple overlapping structures that obscure key features and make identification difficult in 2D. Conceptually, this could be addressed by acquiring data across the entire lamella, which has become feasible due to faster, parallel beam-shift-based tilt series acquisition^14^. However, such strategies are still constrained in practice given that the illuminated area extends beyond the recorded field of view, leading to anisotropic radiation damage from repeated exposures. Recent implementations mitigate this effect by reducing the dose per tilt series but comes at the cost of reduced signal-to-noise ratio, limiting downstream processing and biological interpretation^15^. In addition, such approaches substantially increase microscope time, data processing and storage requirements, as large areas must be recorded at high resolution despite unknown relevance for such detailed analysis beforehand.

Beyond targeting specific ROIs or structures, it is interesting to consider the cellular context surrounding high-resolution volumes for broader integration of cryoET into structural cell biology. In this regard, other approaches such as cryo-FIB-SEM-based volume imaging and soft X-ray tomography give access to extended volumetric context and can be complementary to cryoET^16,17^. However, they require specialised instrumentation and typically operate at lower resolution, on the order of tens of nanometers^16^. In standard cryoET workflows, data acquisition is optimized for high-magnification and target small areas, leaving the surrounding information and most of the lamella unexplored in 3D. As a result, intermediate-scale information on specimen quality and spatial organisation that can be observed during preparation or screening is largely neglected. This creates an opportunity to recover the 3D context directly during cryoET data acquisition itself, without additional hardware or destructive imaging. To address this, we and others have increasingly used low-magnification, high-defocus tomograms to capture so-called mesoscale information^8,18,19^.

Here, we expand on this concept of view tomograms and present a streamlined, semi-automated implementation of mesoscale tomography, coined View Tomo. We show that the acquisition of low-magnification tomograms provides highly relevant information in 3D that complements high-resolution cryoET, with minimal demands on additional instrument time, and data storage or impact on downstream processing, including subvolume averaging. To support this, we developed an integrated workflow and computational tools for robust data acquisition and reconstruction, enabling seamless incorporation of View Tomo, into routine cryoET workflows and specific correlative pipelines. We highlight use cases across a range of biological applications, illustrating how View Tomo enables 3D guided targeting of cellular structures, including the identification of rare membrane remodelling events and viral assembly intermediates. We further show that view tomograms contribute context to downstream high-resolution analysis and enable efficient screening of specimen conditions.

## Results

### View Tomo enables rapid and systematic specimen analysis in 3D

We implemented View Tomo as a semi-automated workflow for rapid and systematic screening of large specimen areas at low magnification, in 3D and in an unbiased manner. The approach complements conventional view-magnification projection images (typically in the range of 2,000-to 8,000-times magnification) by acquiring low-dose, high-defocus tilt series in ~5 minutes using a simple script implemented in SerialEM^20^ (see parameters in Table 1). A step-by-step protocol is provided along with the script as Supplementary Information. The main idea behind View Tomo was to avoid repeated switching between imaging conditions, which implies careful alignment and can be time-consuming, and instead facilitates efficient acquisition of volumetric information across the entire lamellae. The speed of acquisition is primarily achieved by performing tracking directly on the data acquired, and as defocus changes are insignificant at these magnifications, active focusing is not needed. As such, we believe that the workflow is highly valuable but relatively low-key and can be implemented in other interfaces available for data collection without substantial modification.

**Table 1:** Typical View Tomo acquisition parameters. *The range of acquisition times corresponds to tilt series with a total number of 31 to 41 image stacks with 1 s exposure time used per image and illumination set to 50 e^−^px^−2^. **Indicated values show the higher end of expected fluence; interpretable tomograms can be achieved with lower values.

| Magnification | Acquisition time (min)* | Expected Fluence ($\text{e}^{-}\text{\AA}^{-2}$)** | Nyquist frequency ($\text{nm}^{-1}$) | Recommended defocus ($\mu\text{m}$) | Field of view on K3 ($\mu\text{m} \times \mu\text{m}$) |
| --- | --- | --- | --- | --- | --- |
| 2,400x | 4-5 | 1.5 | 8 | 80-150 | 22 x 16 |
| 3,600x | 4-5 | 3.8 | 5 | 40-150 | 14 x 10 |
| 4,800x | 4-5 | 6 | 3.8 | 30-120 | 8 x 5 |
| 8,700x | 4-5 | 16 | 2 | 30-90 | 6 x 4 |

Recommended acquisition parameters based on our experience are given in Table 1. In this context, for cryoET projects where high-resolution subvolume averaging is anticipated, pre-exposure of the sample and electron dose is an important consideration to limit radiation damage. We typically acquire view tilt series (*i*.*e*., the series of projection images collected prior to view tomogram reconstruction) at 2,400-3,600-times magnification, corresponding to total fluences on the order of 1-4 e^−^/Å^2^. At these levels, the additional exposures introduced by View Tomo, in comparison with using only a few view projection images in current workflows, is fully compatible with subsequent high-resolution subvolume averaging^21^. These magnifications are sufficient to resolve the majority of cellular features useful for contextual analysis, such as organelles, cytoskeletal elements and chromatin domains.

In addition to the step-by-step protocol (see Supplementary Information), Figure 1 illustrates the View Tomo workflow. Briefly, after loading the sample into the TEM, a montage of the grid is typically acquired and view magnification tilt series are collected in batch at ROIs for subsequent reconstruction. A key challenge of our approach is reconstructing view tomograms within a timeframe streamlined with microscope sessions and high-resolution cryoET data acquisition. However, existing automated alignment methods were not compatible with view tilt series and did not consistently yield acceptable alignment. This was attributed to the presence of high-contrast features such as edges of the lamella, ice contamination, and vacuum, that dominate cross-correlation during alignment. To address this, we developed a deterministic masking strategy based on histogram matching to suppress these dominant features prior to alignment. This was implemented in a fully automated, single-step reconstruction pipeline utilising AreTomo2^22^ and IMOD Etomo^23^ to reproducibly and quickly yield well-aligned tilt series and interpretable low-magnification tomograms of lamellae with minimal user intervention.

**Figure 1.**
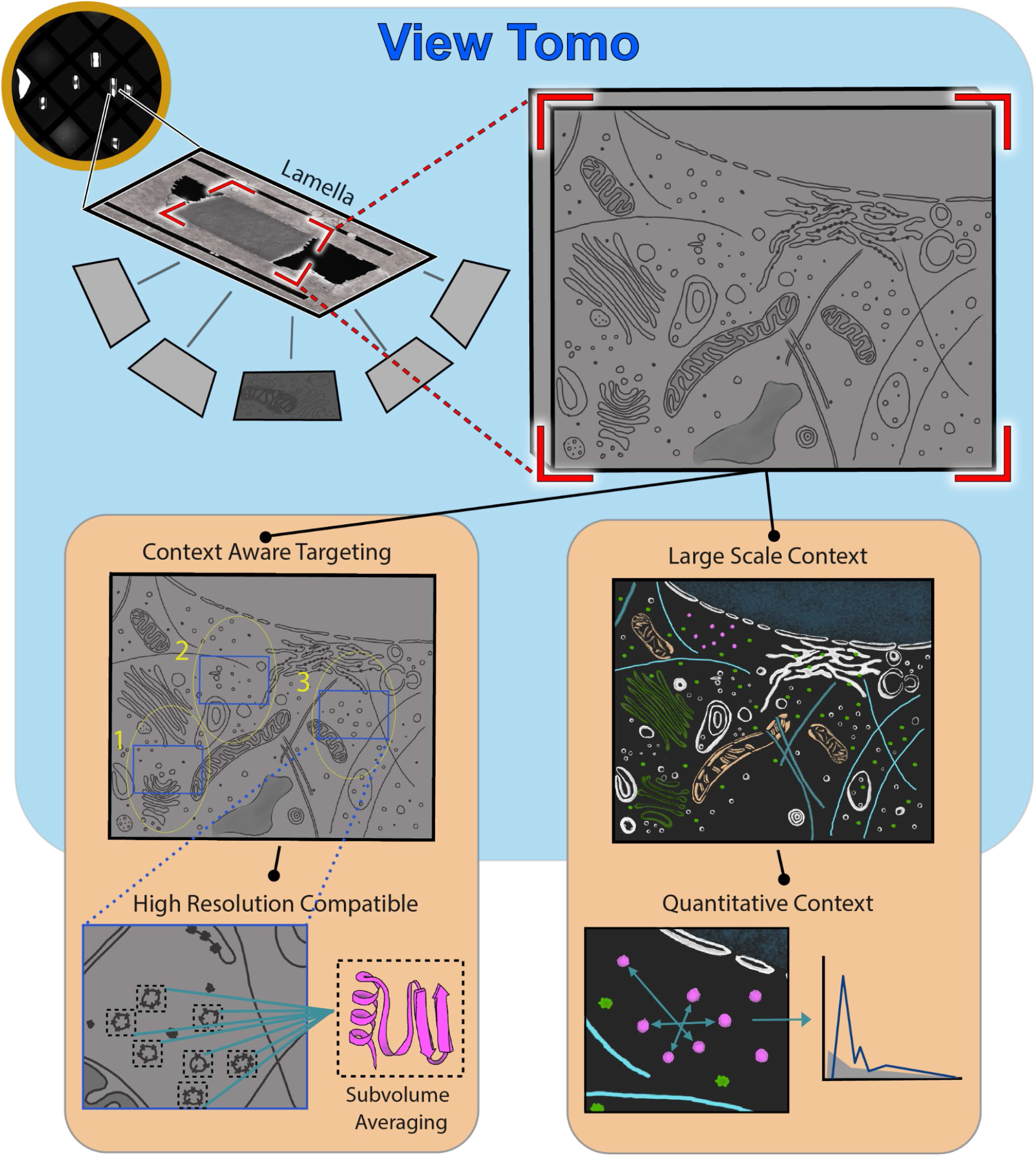
Overview of the View Tomo workflow and its applications. Schematic representation of View Tomo. Low-magnification tilt series are acquired across entire lamellae and reconstructed automatically to generate large-scale tomograms. These volumes are used to guide selection of regions for high-resolution cryoET, while remaining compatible with subsequent detailed analysis and subvolume averaging. In parallel, view tomograms provide information on the organisation of the specimen at larger length scales. Their high contrast supports automated three-dimensional segmentation, enabling identification and quantification of cellular and viral features across the lamella.

This approach supports multiple modes of operation depending on experimental requirements: (i) **real-time acquisition and analysis**, where view tomograms are acquired, reconstructed and interpreted during the session to guide immediate targeting decisions; (ii) **parallel acquisition and analysis**, where multiple users share instrument time, alternating between data collection and analysis; and (iii) **batch acquisition with offline analysis**, where multiple grids and lamellae are mapped using View Tomo, analysed afterwards, and later on revisited for targeted high-resolution data collection. This flexibility makes View Tomo practical across different experimental settings and motivated its application to a range of biological projects and scientific questions in the group and beyond.

#### Case 1: View Tomo enables targeting of membrane assembly intermediates in viral factories

We first applied View Tomo to investigate viral particle assembly during poxvirus infection. VACV represents the prototype of *Poxviridae*, a family of large dsDNA viruses for which viral genome replication and viral progeny assembly take place entirely in the cytoplasm of the eukaryotic host cell, causing extensive remodelling of cellular membranes through the course of infection. Indeed, key viral membrane intermediates with half-moon crescent shape have been reported and arise within densely packed assembly sites, making them difficult to identify in projection images for cryoET. Importantly, assembly is centred around a membrane-driven pathway in which small membrane precursors are recruited to growing membrane crescents with stabilized open ends. These crescents are associated to the scaffold protein D13 imposing curvature on their convex side and expand progressively before closing to form spherical immature virions^24^. Next, several maturation steps by phosphorylation and proteolytic cleavages are necessary to make infectious particles called mature virions with the characteristic brick-like shape^25^.

View Tomo revealed the organisation of VACV assembly sites in 3D, allowing their architecture to be examined directly inside the surrounding cellular environment (Fig. 2a). Notably, these virus-induced compartments, also called viral factories, remained identifiable despite the presence of large ice crystals on the surface of the lamella, which however, rendered individual projection images difficult to interpret (Fig. 2b). Within the crowded and reorganized cytoplasm of VACV-infected cells, view tomograms enabled clear identification of membrane crescents and immature virions but also resolved key intermediates of viral assembly including viral membrane precursors that contribute to crescent growth (Fig. 2). These smaller membrane structures were difficult and nearly impossible to detect in projection images because of low contrast and overlap with neighbouring features. Their visibility in reconstructed volumes is striking and made it possible to target regions enriched in specific stages of viral assembly for subsequent high-resolution cryoET as well as interpretation thereof.

**Figure 2.**
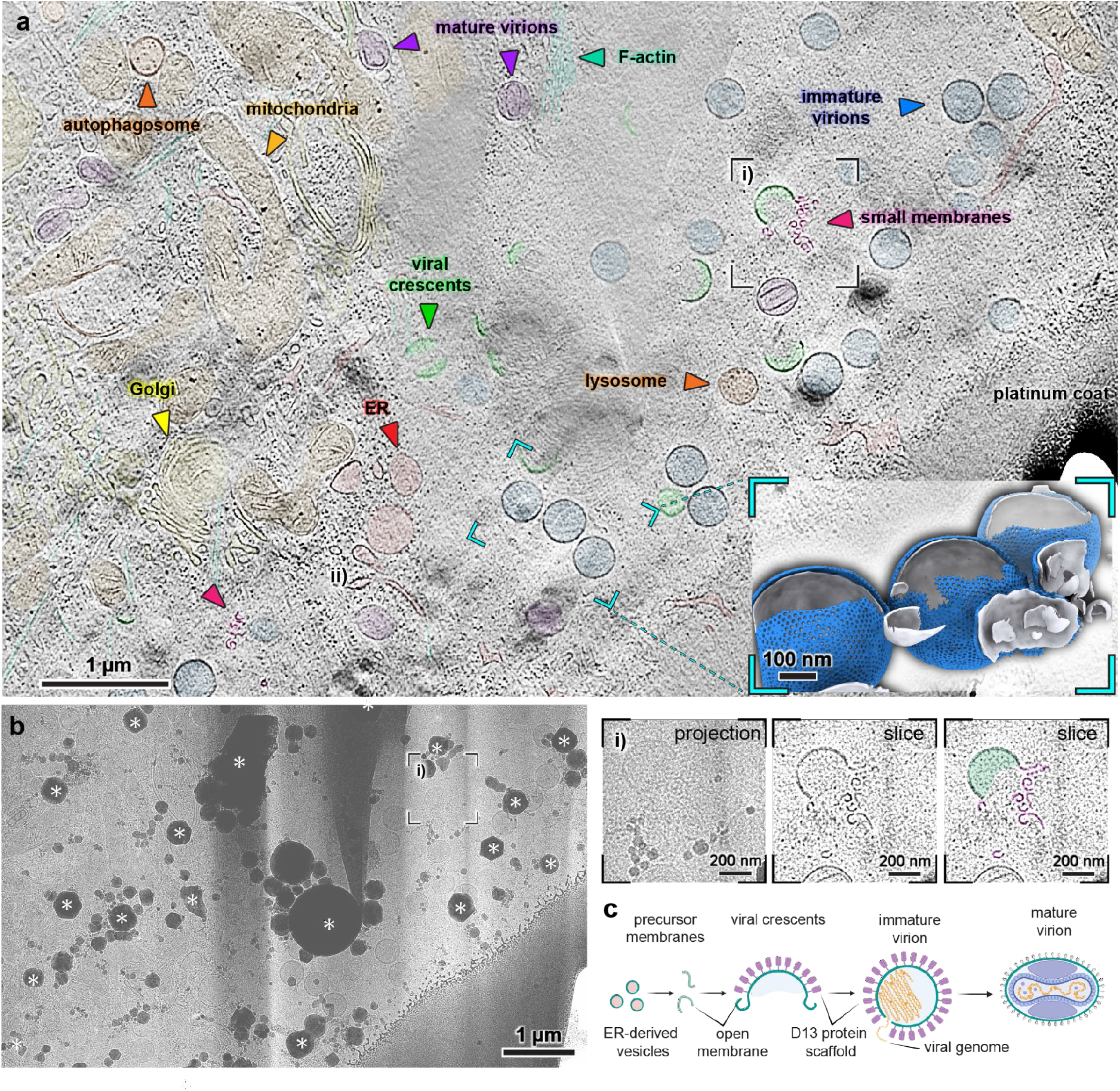
View tomograms guide the localization of specific viral assembly intermediates of VACV in the viral factory. **a**, Slice through a view tomogram (3,600-times magnification) of a HeLa cell 8 h post infection showing a large region of cytoplasm containing assembling viral particles. Major cellular features are annotated: mitochondria (orange), lysosomes and autophagosomes (red–orange), Golgi apparatus (yellow) and ER network (red). Virions at different assembly stages are visible, including viral crescents (green), immature virions (blue) and mature virions (purple). Inset, 3D map derived from a high-magnification tomogram showing viral membranes (grey) and template-matched positions of the D13 scaffold (blue) within the region highlighted in cyan (see Methods). **b**, Left, Projection image of the same lamella, showing reduced interpretability despite visible mitochondria and immature virions. Ice contamination is indicated (*). Right, Enlarged region (i) from **a** and **b**. The viral crescent is difficult to distinguish in the projection image but clearly resolved in the tomographic slice. Nearby small membrane structures are only visible in the tomogram. **c**, Schematic of poxvirus assembly based on VACV, illustrating the intermediates shown in a and b (membranes, green; D13 scaffold, pink; viral DNA, orange).

#### Case 2: View Tomo provides large-volume spatial information complementary to high-resolution cryoET analysis of viral particle assembly

While View Tomo helped identify specific membrane assembly intermediates in VACV directly in the rearranged host cell cytoplasm, we next asked whether the same approach could support both targeted high-resolution imaging and quantitative spatial analysis in a structurally reordered nuclear system upon viral infection. For the human cytomegalovirus (HCMV), a widespread dsDNA virus of the *Herpesviridae* family, capsid assembly and packaging of the viral genome are orchestrated within the host cell nucleus^26^. Using the extended field of view provided by low-magnification tomograms, capsids could be identified directly within the nuclear volume and classified as immature, empty or mature as classically done for herpesviruses (Fig. 3). Such pivotal information was then used to guide targeted acquisition strategies for high-resolution tomograms of specific capsid populations and downstream subvolume averaging.

**Figure 3.**
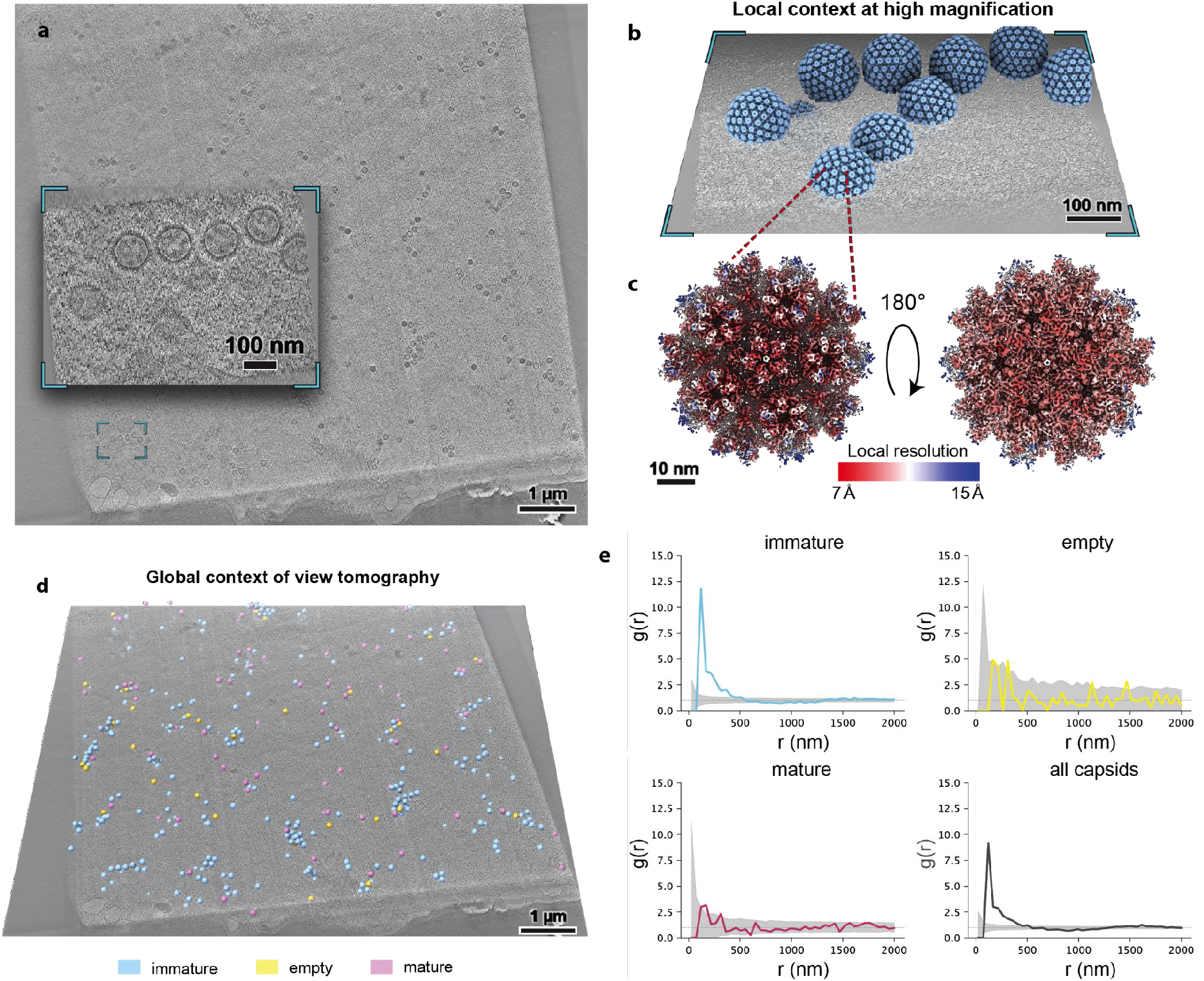
Large-volume contextual information complementary to high-resolution cryoET analysis of HCMV capsid assembly in the nucleus of the infected cell. **a**, A slice through a view tomogram (3,600-times magnification) of HCMV-infected cell nucleus. Inset depicts a slice through a representative high magnification tomogram (64,000-times) on intranuclear HCMV capsids. **b**, High magnification tomogram from **a**. with density maps plotted back into the tomogram coordinate system, which provides local context of HCMV capsid assembly. **c**, Result of subvolume averaging of HCMV penton vertices. The map is colored based on local resolution as indicated. **d**, Slice through the view tomogram from **a**. with capsid particles plotted back and colored based on their type: immature (pale blue), empty (pale yellow), and mature (pale purple). **e**, Spatial distribution analysis of HCMV capsid types in the infected cell nucleus from the view tomogram data in **a**. (see Methods). The radial distribution function g(r), as described by Martinez-Sanchez et al., 2022, quantifies the probability of finding a neighbouring capsid at distance r relative to a random distribution, where g(r) = 1 indicates complete spatial randomness. Functions were calculated for immature, empty, mature and all capsids combined from particle coordinates as in **d**. Grey shading in addition to the plot indicates the 5–95% confidence interval derived from 500 simulations of a null model in which capsids were placed randomly within the segmented nuclear volume. Immature capsids show significant clustering at short range, while mature and empty capsids are distributed indistinguishably from random. HCMV, human cytomegalovirus.

Importantly, the additional exposure introduced by View Tomo did not preclude high-resolution analysis by subvolume averaging of viral capsids (Fig. 3c). Despite the low-magnification pre-exposure, subvolume averaging of the targeted B-capsids (immature capsids) yielded a reconstruction at approximately 9 Å resolution, a regime in which major secondary-structure features can be resolved. These findings highlight that View Tomo can be integrated into high-resolution cryoET workflows for *in situ* structure determination by subvolume averaging.

Furthermore, high magnification data can provide the local context of HCMV capsid assembly and maturation. In parallel, View Tomo provided the global context and enabled a more comprehensive spatial analysis of capsid organisation across the nuclear volume in the infected cell. Classification of capsids directly in the view tomograms allowed mapping of the relative abundance and arrangement of different maturation states within the lamella (Fig. 3d). Preliminary analysis indicated that immature capsids form local clusters, whereas mature capsids appear more dispersed (Fig. 3e). Together, these analyses pave the way for more targeted research to understand how HCMV capsid assembly and maturation are spatially organised within the nucleus. Overall, it also illustrates how View Tomo can reveal significant insights in biology from contextual information and new means for quantitative output of cryoET at different scales.

#### Case 3: View Tomo identifies cellular membrane remodelling events during HSV-1 secondary envelopment

Following egress from the nucleus, herpesviruses undergo additional assembly steps in the host cytoplasm, culminating in acquisition of the viral envelope in a process known as secondary envelopment^27^. At this stage, capsids, tegument components, plus host-derived membranes come together and assemble into mature virions within specialised viral assembly compartments. These compartments form by extensive reorganisation of the host cell at the level of organelles, cytoskeleton and secretory pathway in particular^28^.

We applied View Tomo to examine the organisation of these compartments during secondary envelopment of Herpes Simplex Virus type 1 (HSV-1). A central challenge in these specimens is the coexistence of diverse membrane structures and viral intermediates within a highly crowded and heterogeneous cytoplasmic environment. This complexity is further increased by multiple secondary envelopment pathways reported in the literature such as budding into single vesicles or multiviral bodies^29^.

View tomograms enabled identification of membrane remodelling events associated with HSV-1 secondary envelopment at various stages of virion assembly (Fig. 4, Supplementary Fig. 1). These events are difficult to detect in projection images because membrane contrast is strongly orientation-dependent while the surrounding compartment is structurally complex and heterogenous (Fig. 4b,c). In slices of view tomograms taken on lamellae, however, the sites were evident and could be located directly, enabling targeted acquisition at high-resolution for defined stages of HSV-1 secondary envelopment. Furthermore, viral capsids present in the crowded cytoplasmic environment are difficult to discriminate from small intracellular vesicles based on a single projection image; view tomograms provided the necessary 3D information to confidently allow specific identification of viral capsids (Fig. 4c). Across multiple grids, the collected datasets also enabled systematic mapping of the spatial relationships between viral intermediates and membrane structures within the viral assembly compartment.

**Figure 4:**
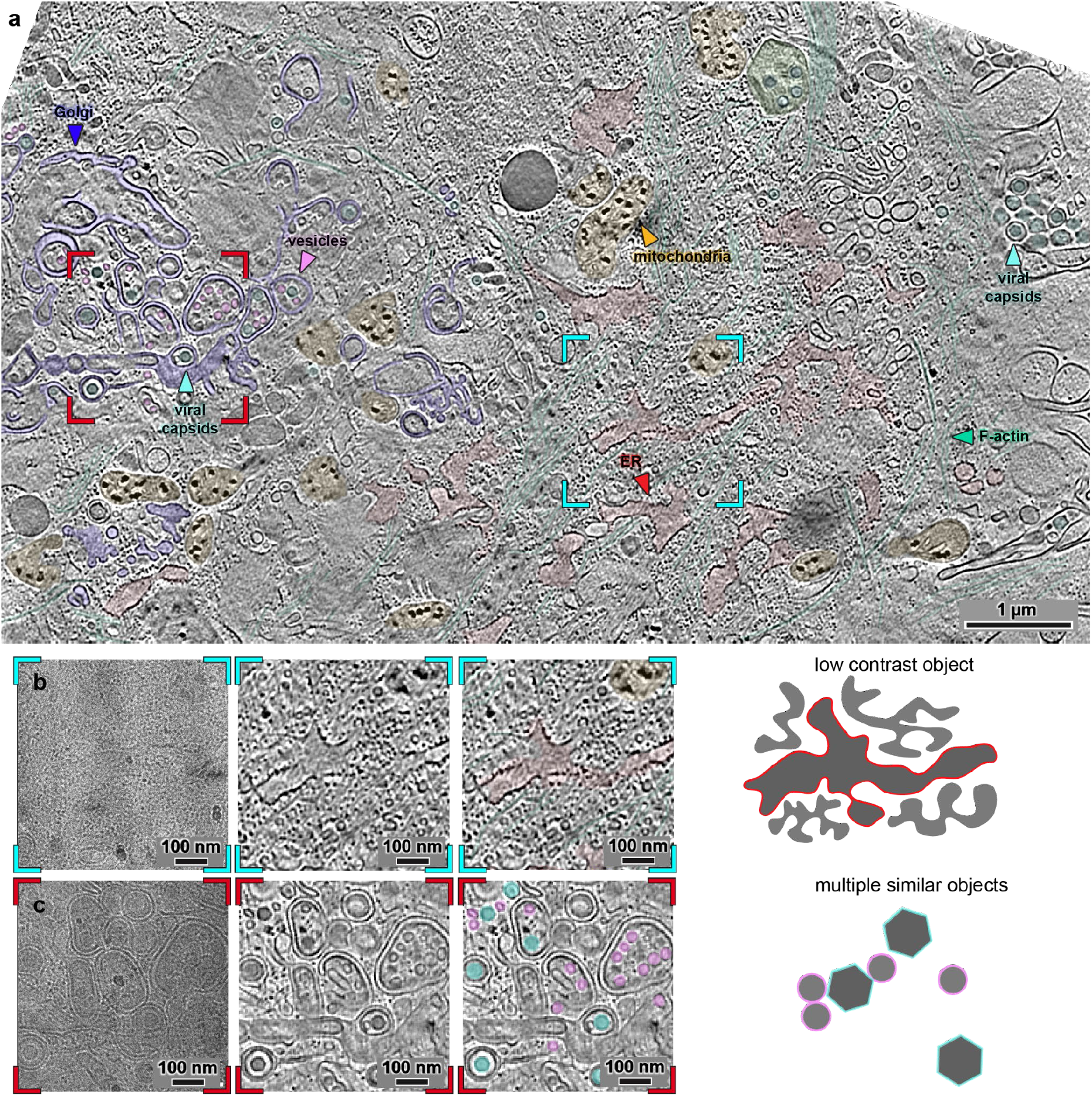
View tomogram-aided target identification of HSV-1 secondary envelopment and associated cellular rearrangements in lamellae. **a**. Slice through a view tomogram at 3,600-times magnification of an HSV-1-infected cell. Coloured brackets indicate magnified areas. b, From left to right: projection image of the region of interest (ROI) in the lamella (left), slice through the corresponding view tomogram, an annotated version of the slice showing ER in red, and a schematic representation of the challenges associated with identifying this target (right). The ER and other membrane-bound organelles are extensively reorganised during HSV-1 infection. In this case, the target structure has low contrast and is almost invisible in the projection image. c, From left to right: projection image of a region containing a viral assembly compartment (left), slice through the corresponding view tomogram, an annotated version with capsids (blue) and vesicles (pink) highlighted (middle/right). Here, the challenge arises from distinguishing objects of interest (capsids) from vesicles of similar size and appearance at view magnification. HSV-1, herpes simplex virus 1; ER, endoplasmic reticulum.

### Other benefits of View Tomo for assessment of sample preparation and automated segmentation of cellular volumes

View Tomo therefore established a framework for more systematic analysis of large-scale cellular organisation during viral infections, as we have seen for large, enveloped dsDNA viruses hereinabove: poxviruses (Fig. 2) and herpesviruses (Fig. 3, 4). In the case of negative-stranded RNA viruses with segmented genomes of the *Phenuiviridae* family, view tomograms of Rift Valley fever virus (RVFV)-infected cells revealed extensive cytoplasmic remodelling, including dense vesicular clusters associated with viral assembly (Fig. 5, Supplementary Fig. 2). The high contrast obtained at low magnification and high defocus enabled automated segmentation of large cellular volumes, allowing membranes, mitochondria, microtubules and viral particles to be identified using a U-net–based approach (see Methods). The contrast is comparable to that of stained plastic sections in conventional EM sample preparation, but without chemical fixation or heavy metal staining^3^. Automated segmentation can be applied on view tomograms and reveal unexpected interplay between various cellular features as well as their spatial reorganization upon infection (Fig. 5). In the case of RVFV, assembly of new viral particles occur inside cytoplasmic viral factories where microtubule spindle-like structures are found adjacent to non-condensed nucleus. Altogether, probing virus-host interactions *in cellula* with View Tomo suggest deviations from the canonical cell cycle organisation and new paradigm in the viral universe.

**Figure 5.**
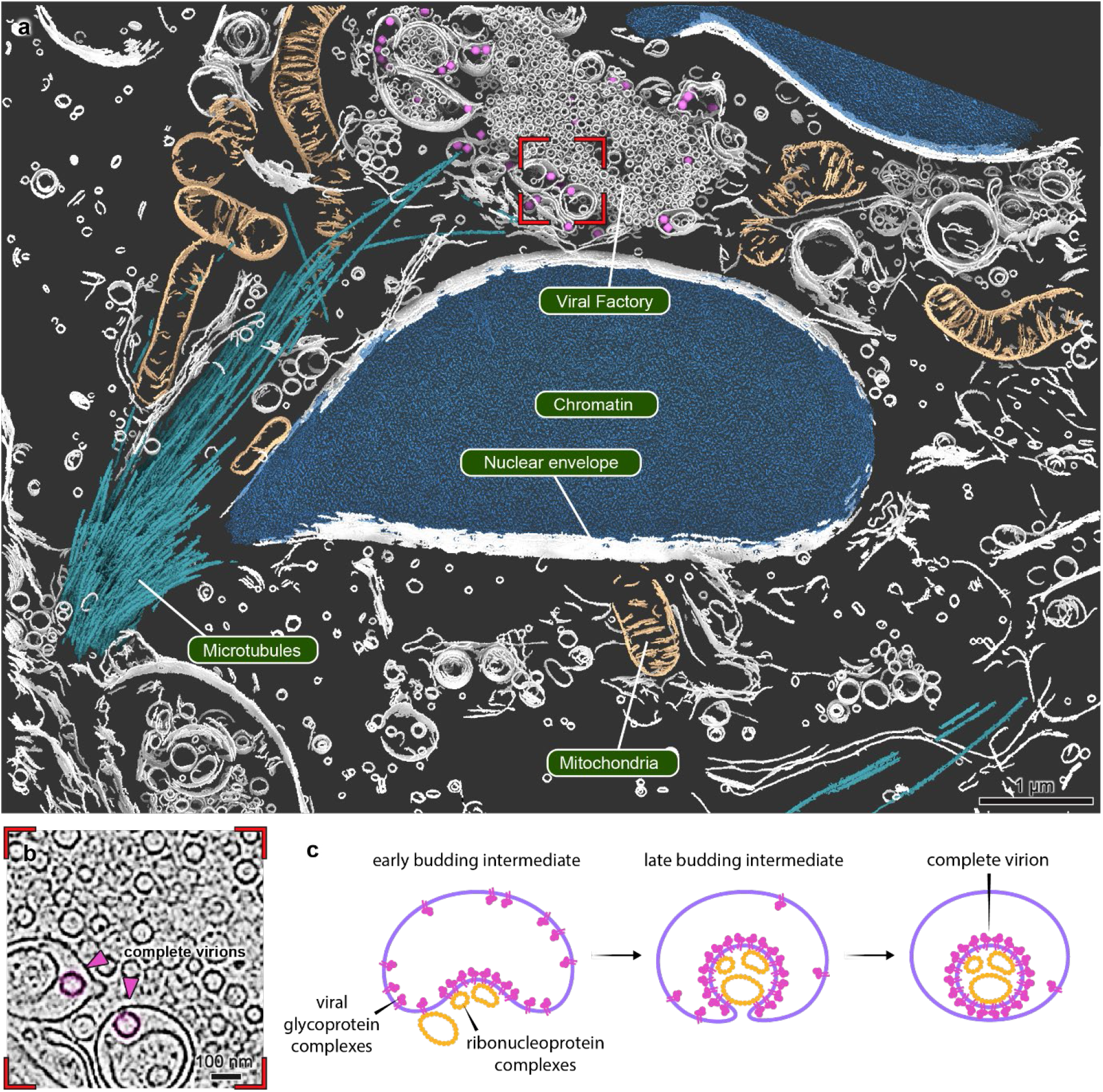
View tomograms enable segmentation of whole cellular compartments. **a**, Three-dimensional segmentation of a view tomogram acquired at 3,600× magnification from a Rift Valley fever virus (RVFV)-infected Vero cell at 18 h post infection (hpi), illustrating large-scale cytoplasmic remodelling during infection. Viral assembly is associated with dense vesicular clusters, and broader changes in cellular organisation are apparent. **b**, Magnified view of the highlighted region in a, showing viral assembly sites and associated membrane structures in detail. **c**, Schematic representation of the RVFV assembly process and the associated reorganisation of the host cytoplasm.

### Additional practical advantages of View Tomo to assess specimen state across preparation conditions

View Tomo was initially motivated by the need to identify specific regions of interest or rare biological events that are difficult to recognise in projection images (Figs. 2–4). While we implemented a robust and automated workflow for acquisition and reconstruction of low-magnification tomograms (Fig. 1), routine use across a wide range of projects revealed additional practical advantages. In particular, view tomograms provide a direct readout of specimen state, both in terms of biological organisation and preservation quality. This is especially valuable for cryoET of cells, tissues and small organisms, where it enables rapid and unbiased assessment of specimen condition, including physiological state and vitrification quality.

This capability is particularly useful during sample preparation and optimisation. View Tomo allowed systematic comparison of experimental conditions such as cell culture conditions or cryo-protectants, and facilitates evaluation of whether relevant biological phenotypes are preserved on the grid (Supplementary Fig. 3). In this way, it supports iterative optimisation and enables more confident selection of regions for subsequent high-resolution data acquisition (Fig. 5).

The data resulting from optimisation experiments may not be suitable for high-resolution structural analysis per se, but they can still support qualitative assessment of specimen state, inform subsequent steps in the workflow and enable hypothesis generation that may lead to new biological discoveries (Supplementary Fig. 4). Taken together, these observations show that View Tomo extends cryoET workflows beyond targeting alone, enabling more efficient experimental design and integration with correlative imaging approaches while remaining compatible with downstream structural analysis.

### Towards integration with (super-resolution) correlative workflows and post-TEM cryo-fluorescence imaging

#### 1. Guiding high-resolution data acquisition

In addition to assessing specimen quality, View Tomo provides a practical guide for subsequent high-resolution imaging. The reconstructed volumes define the usable tilt range and determine ROIs that remain unobstructed across tilts for example. In combination with correlative workflows, it enables more reliable selection of acquisition areas and regions used for tracking and focusing. This reduces trial-and-error during high-magnification data collection on high-end microscopes and enables more confident targeting of phenotypes or ROIs. Overall, all these characteristics of View Tomo increase the throughput of high-quality datasets, simplifying downstream processing and reducing computational overhead.

#### 2. Integration with post-TEM cryo-fluorescence imaging

In conventional cryoCLEM workflows, fluorescence imaging is typically performed prior to observations at the TEM, given that radiation damage from beam exposure during high-magnification tilt series acquisition can compromise the fluorescent signal. As super-resolution cryoCLEM becomes more widely accessible, the increased acquisition time required by such techniques will result in growing demands on cryo-fluorescence microscopy in terms of machine time and data analysis, making efficient use of imaging time increasingly important at this stage as well.

Building on observations that low-magnification TEM imaging does not impair subsequent cryo-fluorescence measurements^30^, we explored the feasibility of performing super-resolution cryo-fluorescence imaging after View Tomo data acquisition. In this workflow, view tomograms are acquired at a magnification that closely matches the spatial scale of super-resolution cryo-fluorescence imaging, enabling direct comparison between the two modalities. In our implementation, view tilt series acquired at ~8,700× magnification remained compatible with subsequent cryo-fluorescence imaging, yielding localisation precision on the order of ~20 nm (Fig. 6, Supplementary Fig. 5). Depending on the electron fluence used, this approach may also remain compatible with subsequent reloading into the TEM for targeted high-resolution data acquisition. While a detailed characterisation of this workflow is presented elsewhere^31^, these observations suggest that View Tomo can help decouple region selection from fluorescence acquisition, enabling more efficient use of time-intensive correlative imaging approaches.

**Figure 6.**
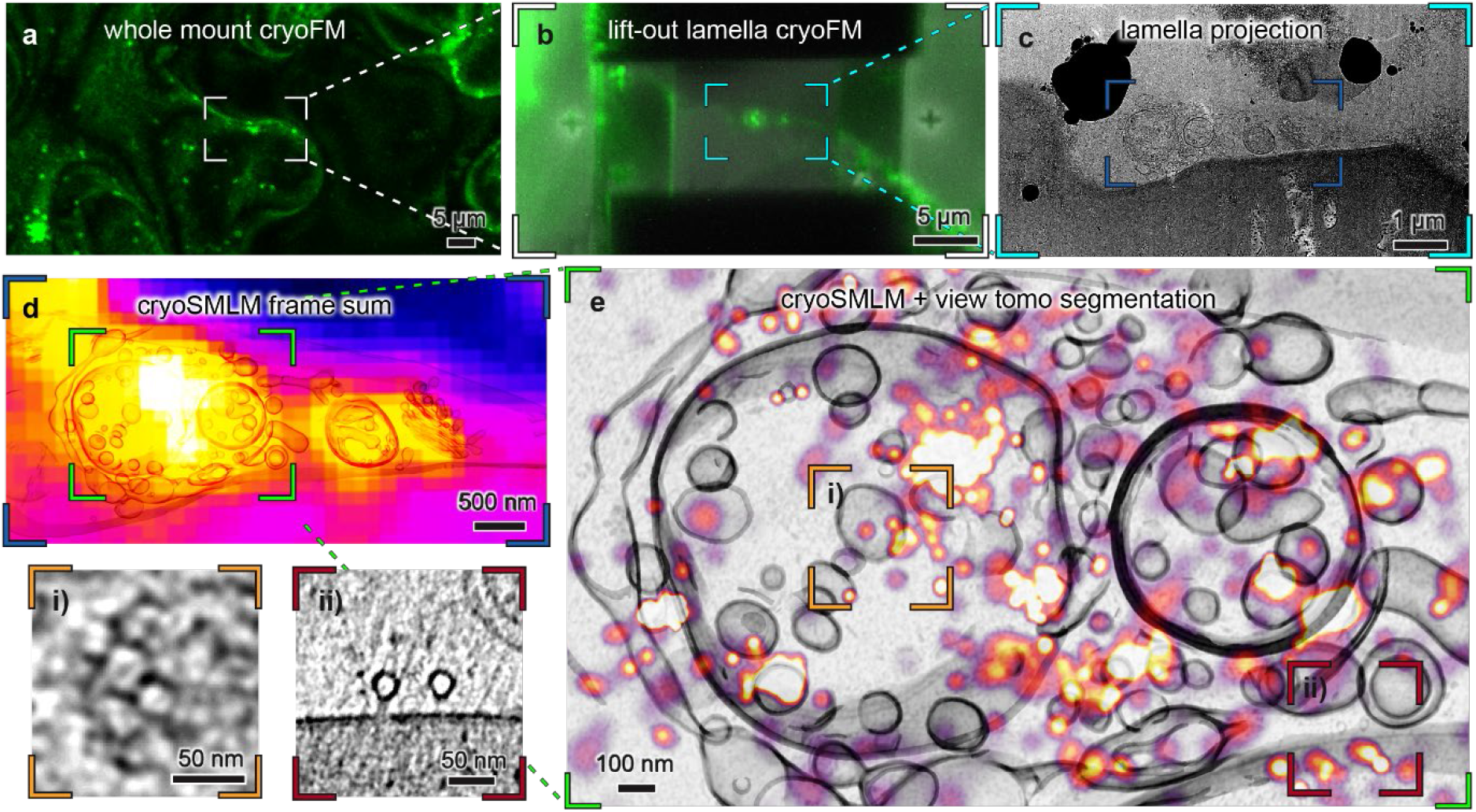
Post-TEM super-resolution cryoCLEM following View Tomo. **a-c**, Correlative cryoCLEM workflow for a *Nicotiana benthamiana* leaf expressing ATG9:GFP. The sample was high-pressure frozen and a lift-out lamella was prepared by cryoFIB-SEM. **d**, Following acquisition of a view tomogram at 8,700-times magnification, the lamella was transferred to a cryo-fluorescence microscope equipped with an ultrastable cryogenic stage for cryo-single molecule localisation microscopy (cryoSMLM) data acquisition. The fluorescence signal acquired after TEM imaging is comparable to that observed prior to TEM (iFLM, Supplementary Fig. 5). The panel shows an overlay of the sum of the cryoSMLM data and the cryoET data. **e**, Overlay of a projection of the segmented membrane volume from the region shown in a–d with the corresponding cryoSMLM. The majority of ATG9:GFP localisations overlap with small membrane vesicles observed inside or around the two autophagosome-like structures. **i-ii**, Magnified regions indicated in e showing sections through the view tomogram. i, Detail of a clathrin cage within the autophagosome-like compartment. ii, Projection along two microtubules with associated F-actin filaments adjacent to the plasma membrane.

## Discussion

View Tomo extends cryoET workflows by improving the identification and selection of biologically relevant targets. For *in situ* cryoET, the combination of thin lamellae and highly specific cellular states means that even common processes become effectively rare within the imaged volume. Screening based on 2D projection images is therefore intrinsically limited, as overlap of cellular components and orientation-dependent visibility obscure many relevant features, particularly membranes and pleomorphic structures. By shifting screening from projections to reconstructed volumes, View Tomo increases the likelihood that high-resolution datasets are collected from regions containing the appropriate biological context, reducing both missed targets and uninformative acquisitions. Although we focus here on FIB-milled lamellae, the underlying approach is broadly applicable and can be extended to a wide range of specimen types in the TEM.

Beyond targeting, View Tomo provides a reusable three-dimensional record of the specimen at the scale of the entire lamella. This enables high-resolution datasets to be interpreted within their surrounding cellular environment, rather than as isolated subvolumes, and supports additional modes of analysis based on spatial organisation. In particular, the availability of mesoscale 3D information from the same specimen volume opens the possibility of quantitative approaches, such as spatial or point-pattern analysis, that complement high-resolution structural data. View tomograms constitute not only a screening tool but a complementary dataset that links molecular structures to their cellular environment.

Finally, View Tomo extends the practical utility of specimens that are not suitable for high-resolution cryoET. While non-vitreous or otherwise suboptimally preserved samples cannot support reliable high-resolution structural interpretation, biological interpretation is routinely performed in conventional electron microscopy under preservation criteria that differ from those of vitrified cryo-samples, i.e. the lack of gross freezing artefacts. Low-magnification tomograms therefore provide useful readouts of specimen state, spatial relationships and large-scale cellular architecture, supporting sample optimisation and hypothesis generation when high-resolution data collection would not be justified. More broadly, because such datasets can be acquired at low total fluence, this approach may be compatible with subsequent correlative workflows, including super-resolution cryo-fluorescence microscopy, enabling iterative imaging strategies that combine contextual ultrastructure with molecular specificity.

## Materials and Methods

### Mammalian cell culture

HFF-hTERT and HeLa cells were both cultured in Dulbecco's modified Eagle's medium (DMEM) supplemented with 1x GlutaMAX (Gibco) and 10% (v/v) fetal bovine serum (FBS). Vero cells were cultured in DMEM supplemented with 1x GlutaMAX (Gibco) and 5% UltraGRO (human platelet lysate, hPL). All cells were cultured at 37°C with 5% CO_2_. Before infection the cells were maintaed at 80% confluency.

### VACV infection on grids

Holey SiO_2_ film on R 1/4 gold 200 mesh grids were glow discharged for 90 s at negative polarity and 25 mA plasma current (Quorum GloQube Plus). Grids were coated with fibronectin solution (50 µg/ml in 20 mM HEPES) for 2 h at 37 °C to promote cell adhesion. HeLa cells were seeded at a cell density of 150 cells mm^−2^ and grown in complete medium (DMEM + 10% (v/v) FBS + 1x GlutaMAX) overnight. Cells were infected with VACV (3.24 × 10^8^ PFU/ml stock from J. Krijnse-Locker, Paul-Ehrlich-Insitut, Langen, Germany) at an multiplicity of infection (MOI) of 10 for 30 min at room temperature and 15 min at 37°C. The viral inoculum was replaced with complete medium and infection was allowed to progress for 8 h at 37°C with 5% CO_2_. For the last 30 min, 5 μg mL^−1^ Hoechst 33342 DNA dye was added to the growth media. The VACV infected samples were plunge-frozen using a Leica EM GP2 instrument (chamber temperature 37°C, 80 % relative humidity, blotting time 6-8 s) and clipped into FIB-optimized autogrids. Lamellae were prepared in an Arctis plasma-FIB-SEM instrument operated via WebUI software. Target positions for lamella milling were identified based on the fluorescence signal of Hoechst-stained extranuclear DNA in the viral assembly compartment as detected in the blue channel of the Arctis in-line fluorescence microscope. Lamellae were milled in automation and polished manually to a final thickness of 150-300 nm.

### HCMV infection on grids

Holey SiO_2_ film on R 1/4 gold 200 mesh grids were glow discharged for 45 s at negative polarity and 25 mA plasma current (Quorum GloQube Plus). Grids were placed into tan ibidi µ-slide 2-well co-culture chamber wells and coated with fibronectin solution (50 µg/ml in PBS) for 1 hour at 37 °C to promote cell adhesion. HFF-hTERT cells were seeded at a cell density of 3000 cells per well and grown in complete medium (DMEM + 10% (v/v) FBS + 1x GlutaMAX) for 4 h. Cells were infected with HCMV (TB40/E) mNeongreen-UL112-113 (produced in HFF-hTERT at concentration of 3 × 10^7^ PFU ml^−1^) at an MOI of 30 in serum-free media for 1 h at 37°C. The viral inoculum was replaced with complete medium and infection was allowed to progress for 120 h at 37°C with 5% CO_2_. The HCMV infected samples were blotted for 5 s from the back side of the grid (chamber temperature 37 °C and 50% relative humidity) and plunge-frozen using a Leica EM GP2 instrument and clipped into the FIB-optimized autogrids. Lamellae were prepared on an Arctis plasma-FIB-SEM (Thermo). The milling was guided by the presence of mNeongreen signal in the cell nucleus. Grids were GIS coated and sputter coated, before milling. Lamellae were thinned to approximately 200 nm using an automated procedure. Lastly, a thin final sputter coat was applied before unloading.

### HSV-1 infected cells on grids

Holey SiO_2_ film on R 1/4 gold 200 mesh grids were glow discharged for 60 s at negative polarity and 25 mA plasma current (Quorum GloQube Plus). Grids were coating with fibronectin solution (50 µg/ml in 20 mM HEPES) for 30 min at 37 °C to promote cell adhesion. HFF-hTERT cells were seeded at a cell density of 100 cells/ grid and grown in complete medium (DMEM + 10% (v/v) FBS + 1x GlutaMAX) for 16 h. Cells were infected with HSV-1 mCherry-gC (strain 17+) at an MOI of 10 in serum-free media for 1 h at 37°C. The viral inoculum was replaced with complete medium and infection was allowed to progress for 16 h at 37°C with 5% CO_2_. The grids were the blotted from the back of the grid for 8 s using a Leica EM GP2 instrument at 37 °C and 90% relative chamber humidity, plunge-frozen and clipped into the FIB-optimized autogrids. Lamellae were milled semi-automatically at an Aquilos cryo-FIB-SEM with built in LM module (Thermo) using AutoTEM software. Target cells were chosen based on the fluorescence signal. Final polishing was performed manually until the thickness of approximately 150 nm was reached.

### RVFV infection on grids

As previously described^32^, holey SiO_2_ film on R 1/4 gold 200 mesh grids were glow discharged for 30 s at negative polarity and 25 mA plasma current (Quorum GloQube Plus). Grids were coated with fibronectin by incubation face down on a 20 µL droplet of fibronectin solution (50 µg/ml) for 30 min. Grids were washed briefly in cell culture medium and transferred into wells of an ibidi µ-slide 2-well co-culture chamber prefilled with 60 µl medium. Approximately 200 cells were seeded onto the central region of each grid and allowed to adhere for 3 h. For infection with RVFV, cell culture medium was exchanged with 50 µl Rift Valley fever virus (strain MP-12) inoculum at a concentration of 1 × 10^7^ PFU/ml per grid, corresponding to an MOI above 1. The RVFV stock had been produced in Vero cells maintained in DMEM supplemented with 1× GlutaMAX and 2% FBS, and the inoculum therefore contained 2% FBS. Cells were incubated with viral inoculum for 18 h at 37°C with 5% CO_2_. Cells were vitrified using a manual plunge freezer custom-built in the Department of Biochemistry at the Max Planck Institute, Martinsried, Germany. Lamellae were produced using automated procedure in an Arctis plasma-FIB-SEM (Thermo).

### View Tomo data acquisition and processing

TEM data were collected using SerialEM software on a Titan Krios microscope operated at an acceleration voltage of 300 kV and equipped with a BioQuantum energy filter set with a slit width of 20 eV inserted and a Gatan K3 direct detector.

View tomograms were acquired using the script presented in this manuscript at nominal magnifications of 2250-times and 3,600-times. Acquisition was performed bi-directionally from 0° to ±51° using 3° increments. For the view tomogram presented in the results the acquisition parameters are as follows:

| **Figure in Results** | **Pixel Spacing (Å)** | **Exposure Time (s)** | **Magnification times (-)** | **Z size (tilts)** |
| --- | --- | --- | --- | --- |
| Figure 2 (VACV) | 24.09 | 1.00134 | 3600 | 35 |
| Figure 3 (HCMV) | 24.09 | 1.00134 | 3600 | 35 |
| Figure 4 (HSV-1) | 23.95 | 1.00134 | 3600 | 35 |
| Figure 5 (RVFV) | 24.09 | 2.00267 | 3600 | 34 |
| Figure 6 (*Nicotiana*) | 38.68 | 2.00267 | 2250 | 43 |

After data collection, raw view tilt series were sorted according to increasing tilt angles using newstack and the reordered stacks used as input for tomogram reconstruction. View tilt series alignment was performed using *view tomo_align*, Aretomo2 or using fiducial tracking in IMOD/Etomo. High-magnification tilt series were aligned either by marker-free alignment in AreTomo2 or by fiducial-based procedures, depending on the dataset. Aligned stacks were ctf-corrected using the estimation from IMOD’s ctfplotter and filtered during reconstruction with a SIRT-like filter equivalent to 15 iterations. Reconstructed 3D-volumes were further band-pass-filtered using mtffilter. For visualisation purposes, stripes resulting from uneven lamella surfaces were removed using *lonardo_destripe*^33^.

### VACV *high-resolution tomogram collection, processing*, D13 plotback and segmentation

High-magnification tilt series were acquired at 42,000-times magnification with a calibrated pixel size of 2.121 Å pixel^−1^ using a dose-symmetric scheme from −60° to +60° with 3° increments using PACEtomo. Tilt series were initially aligned based on patch tracking using IMOD and reconstructed at a pixel size of 10 Å for analysis. Selected tilt series were aligned using metal particles embedded into the lamella after milling and other features as fiducial markers in IMOD.

Membranes of IVs decorated with the scaffold protein D13 were manually segmented in IMOD and a model with initial orientations for D13 created based on custom scripts based on open3d. For the initial average, approximately 500 D13 hexamers were picked manually and averaged in PEET^34^. IV segmentations were used to do template matching with the D13 average as reference in PEET and results cleaned based on cross-correlation score. A six fold symmetrized D13 average was calculated and used as volume for the plotback of D13 into the tomogram coordinate system based on positions and orientations of particles after alignment using a custom python script.

Automatic membrane segmentations for visualization purposes were created using membrain-seg^35^ based on the pre-trained segmentation model checkpoints v10_alpha available from the authors’ GitHub page (https://github.com/teamtomo/membrain-seg) and manually cleaned in ChimeraX^36^.

### HCMV high-resolution tilt series collection, processing and subvolume averaging

High-magnification tilt series were acquired at 64,000-times magnification with a calibrated pixel size of 1.363 Å px^−1^ using a dose-symmetric scheme from −40° to +40° with 2° increments using PACEtomo. Tilt series were aligned using metal particles embedded into the lamella after milling as fiducials. For the initial average, capsid centres were picked manually and averaged against an EMD-18480 map lowpass filtered to 60 Å as a reference in PEET. Penton positions were extracted using custom scripts based on TEMPy. The map was then unbinned and refined at binning 2, reaching a resolution of 14 Å. Using custom scripts, PEET files were converted into RELION-compatible star files, and tilt-series alignments and particle coordinates were imported into WarpTools. Frames were re-aligned using WarpTools. Particles were re-extracted as 2D stacks in Warp, preserving the orientations determined in PEET, and further refined in RELION 5 and M. Final M-refinement with fivefold symmetry yielded a reconstruction at 9.1 Å resolution.

### HCMV capsid classification and spatial clustering analysis

The low-magnification view tomogram was filtered (SIRT-like filter 12) and capsids were picked manually into three IMOD model objects based on their content: scaffold-containing, empty, or DNA-filled. Particles were visualised in ChimeraX overlaid onto the view tomogram. The IMOD model file was converted into a coordinate list using the *model2point* command. To calculate the radial distribution function g(r)^37^ for each capsid population, pairwise distances between all particles of the same type were computed and binned into radial shells. A null model was generated by randomly placing the same number of particles within the same volume. g(r) was obtained by normalising the observed binned distance distribution by the mean of 500 null model simulations, with the 5-95% confidence interval derived from the distribution across simulations.

### Segmentation of RVFV infected cell

The tomogram at binning 3 was loaded into the Dragonfly (Comet) software and the grey scale was normalised. The features were first manually segmented using local otsu brush in small bounding box. Every desired feature was segmented as a separate class; unwanted features were classified as background. The model was trained in 2.5D mode and validated using the whole tomogram. The segmentation was refined and loaded into ChimeraX for manual cleaning, colouring and visualization.

### Nicotiana benthamiana samples Plant growth and cultivation

*Nicotiana benthamiana* plants (wild type and transgenic lines) were cultivated in a growth chamber at 24°C. Plants were grown in a substrate mixture consisting of organic soil in a 3:1 ratio with sand and Sinclair’s 2-5 mm vermiculite. Plants were maintained under a 16 h light / 8 h dark photoperiod and used at 4-5 weeks of age.

### High-pressure freezing

*N. benthamiana* leaves at 4 days post infection, expressing ATG9:GFP, were vacuum infiltrated with 40% Ficol-70, 180 mM glycerol and 20 mM MES pH 5.5 approximately 1h before freezing. Samples were frozen in an HPM010 high-pressure freezer (Baltec) using the 200 µm recess of a type-A carrier paired with the flat side of a type-B carrier (Wohlwend).

### Cryogenic fluorescence screening and serial lift-out

Carriers containing high-pressure frozen leaves were loaded into either a Stellaris 5 microscope (Leica) equipped with a cryogenic stage. Confocal Z-stacks were acquired at 1 µ increments. The carriers were then transferred to an Aquilos 2 dual-beam microscope (Thermo Fisher Scientific) equipped with an integrated fluorescence microscope (iFLM).

Serial lift-out was performed as described previously^18,38^. After platinum sputtering and organoplatinum coating using a GIS, a block of approximately 60 × 25 × 15 µm was milled with a 16 nA probe. Attachment was performed using a single-pass regular cross-section pattern at 0.3-1 nA. The block was attached to a lift-out needle via a copper block, lifted from the carrier, and transferred to a copper 400 × 100 mesh grid (Agar Scientific). It was positioned over two grid bars, welded in place, and sectioned into 2-3 µm slices using a line cut. Slices were trimmed to remove material in front of and behind regions of interest identified using iFLM and milled to ~200 nm thickness.

### CryoSMLM

CryoSMLM data was acquired with the VULCROM setup described in Falckenhayn *et al*.^*31*^. A 488 nm laser with an intensity of 200 W/cm^2^ was used for fluorescence excitation and stimulation of photo-blinking. Raw data was recorded with an EMCCD camera (iXon Ultra 888, Andor) with exposure time of 50 ms and an electron multiplying gain of 700 over the course 1.4 h. Data was binned to 500 ms to increase the signal-to-noise ratio for the cryoSMLM reconstruction. CryoSMLM reconstructions were done as described previously^31^. Transformation of coordinate systems for correlation of cryoSMLM with cryoET was done as described in Pražák *et al*.^*30*^. Specifically, crosses that have been milled into the lamella were used as reference points for the coordinate transformation. The Control Point Selection Tool of MATLAB (Mathworks) was used to determine the coordinate transformation between cryoSMLM and cryoET.

## Acknowledgements

We are grateful to Prof. Lindsay Baker (Department of Biochemistry, University of Oxford) for valuable discussions, and to Dr. Susan Black (Careers Service, University of Oxford) for reviewing the manuscript. Part of this work was performed at the CryoEM multi-user Facility at Centre for Structural System Biology (CSSB), headed by Kay Grünewald and supported by the UHH and DFG (grants INST 152/772-1, 774-1, 775-1 and 777-1). We also acknowledge excellent support by Carolin Seuring, Cornelia Cazey, and Ulrike Laugks, for sample preparation and data acquisition, and Wolfgang Lugmayr for supporting workflows for cryoEM/ET data processing on the Maxwell compute cluster. Portions of this work were performed at Oxford Particle Imaging Centre (OPIC) electron microscopy facility, which was founded by a Wellcome JIF award (060208/Z/00/Z), and Central Oxford Structural Molecular Imaging Centre (COSMIC). We are grateful for the excellent support of Loic Carrique (OPIC), Helen Duyvesteyn (OPIC), Thomas Walther (OPIC), Rishi Matadeen (COSMIC) and Flavia Moreira-Leite (COSMIC), and for training provided by Pranav Shah (STRUBI, University of Oxford). Rift Valley fever virus strain MP-12 was kindly provided by the Friedrich-Loeffler-Institut, Germany. Work on Rift Valley fever virus was funded by the Deutsche Forschungsgemeinschaft (DFG, German Research Foundation) – 518614462; and SFB 1648/1 2024 – 512741711. E.Q. was awarded a Klaus Tschira Boost Fund in 2021 and received funding from the ATIP-Avenir young investigator programme of the CNRS in 2022. M.Z. holds a PhD scholarship from the CEA-AID program and is supported by core funding from Synchrotron SOLEIL. J.F. and R.K. are funded by a Volkswagen Foundation Freigeist Fellowship to R.K. (91671; 91671-2). E.A.M. received Postdoctoral Fellowship from the European Union’s Horizon research and innovation programme under the Marie Skłodowska-Curie grant agreement No 101065943. Enrico Caragliano was funded by the Deutsche Forschungsgemeinschaft (DFG, German Research Foundation) under Germany’s Excellence Strategy EXC 2155 project no. 390874280 and through the DFG Research Unit FOR5200 DEEP-DV (443644894) project BO 4158/5-1 and BO 4158/5-2.

## Authors and Affiliations

Centre for Structural Systems Biology, 22607 Hamburg, Germany

Robert Gebauer, Yuliia Mironova, Märit-Runa Jönsson, Jedrzej Mazur, Clara Feldmann, Enrico Caragliano, Julian Falckenhayn, Jan Hellert, Rainer Kaufmann, Kay Grünewald, Vojtěch Pražák

Leibniz-Institut für Virologie, 20251 Hamburg, Germany

Robert Gebauer, Yuliia Mironova, Jędrzej Mazur, Clara Feldmann, Enrico Caragliano, Jan

Hellert, Kay Grünewald, Vojtěch Pražák

Department of Biochemistry, University of Oxford, Oxford OX1 3QU, UK

Emily Machala, Emma Silvester, Vojtěch Pražák

Department of Life Sciences, Imperial College London, London SW7 2AZ, UK

Enoch Lok Him Yuen, Tarhan Ibrahim, Tolga O. Bozkurt

Oxford Particle Imaging Centre, Division of Structural Biology, Centre for Human Genetics,

University of Oxford, Oxford, UK

Kay Grünewald

Department of Chemistry, University of Hamburg, 20146 Hamburg, Germany

Märit-Runa Jönsson, Kay Grünewald

Department of Physics, University of Hamburg, 22607Hamburg, Germany

Julian Falckenhayn, Rainer Kaufmann

Electron Bio-imaging Centre (eBIC), Diamond Light Source, Harwell Science and Innovation

Campus, Didcot, OX11 0DE, UK

Vojtěch Pražák

Université Paris-Saclay, CEA, CNRS, Institute for Integrative Biology of the Cell (I2BC), 91190 Gif-sur-Yvette, France

Emmanuelle Quemin, Marcel Zimmeck

Synchrotron SOLEIL, L’Orme des Merisiers, 91190 Saint-Aubin, France Marcel Zimmeck

Department of Cell Biology and Infection, Institut Pasteur, 25-28 rue du Docteur Roux, 75015 Paris, France: current address of Clara Feldmann

Hannover Medical School, Institute of Virology, Hanover, Germany

Enrico Caragliano

Cluster of Excellence RESIST (EXC 2155), Hannover Medical School, Hannover

Enrico Caragliano

## References

1 Baker, L. A., Grange, M. & Grünewald, K. Electron cryo-tomography captures macromolecular complexes in native environments.

2 Baumeister, W. Cryo-electron tomography: A long journey to the inner space of cells. Cell 185, 2649–2652 (2022). 10.1016/j.cell.2022.06.034

3 Quemin, E. R. J. et al. Cellular Electron Cryo-Tomography to Study Virus-Host Interactions. Annu Rev Virol 7, 239–262 (2020). 10.1146/annurev-virology-021920-115935

4 Liedtke, J., Depelteau, J. S. & Briegel, A. How advances in cryo-electron tomography have contributed to our current view of bacterial cell biology. J Struct Biol X 6, 100065 (2022). 10.1016/j.yjsbx.2022.100065

5 Schiøtz, O. H., Klumpe, S., Plitzko, J. M. & Kaiser, C. J. O. Cryo-electron tomography: en route to the molecular anatomy of organisms and tissues. Biochem Soc Trans 52, 2415–2425 (2024). 10.1042/bst20240173

6 Nogales, E. & Mahamid, J. Bridging structural and cell biology with cryo-electron microscopy. Nature 628, 47–56 (2024). 10.1038/s41586-024-07198-2

7 Villa, E., Schaffer M Fau - Plitzko, J. M., Plitzko Jm Fau - Baumeister, W. & Baumeister, W. Opening windows into the cell: focused-ion-beam milling for cryo-electron tomography.

8 Moser, F. et al. Cryo-SOFI enabling low-dose super-resolution correlative light and electron cryo-microscopy. Proceedings of the National Academy of Sciences of the United States of America (2019). 10.1073/pnas.1810690116

9 Klumpe, S. et al. A modular platform for automated cryo-FIB workflows. eLife 10, e70506 (2021). 10.7554/eLife.70506

10 Wolff, G., Hagen, C., Grünewald, K. & Kaufmann, R. Towards correlative super-resolution fluorescence and electron cryo-microscopy. Biology of the Cell 108, 245–258 (2016). 10.1111/boc.201600008

11 Dahlberg, P. D. & Moerner, W. E. Cryogenic Super-Resolution Fluorescence and Electron Microscopy Correlated at the Nanoscale. Annual Review of Physical Chemistry 72, 253–278 (2021). 10.1146/annurev-physchem-090319-051546

12 DeRosier, D. J. Where in the cell is my protein? Quarterly Reviews of Biophysics 54, e9 (2021). 10.1017/S003358352100007X

13 Mazal, H., Wieser, F. F. & Sandoghdar, V. Insights into protein structure using cryogenic light microscopy. Biochem Soc Trans 51, 2041–2059 (2023). 10.1042/bst20221246

14 Eisenstein, F. et al. Parallel cryo electron tomography on in situ lamellae. Nat Methods 20, 131–138 (2023). 10.1038/s41592-022-01690-1

15 Hylton, R., Boiero Sanders, M., Prajica, A., Rice, G. & Raunser, S. Streamlined Montage Cryo-Electron Tomography for Exploring the Ultrastructure of Cells and Tissues. bioRxiv, 2025.2009.2001.673430 (2025). 10.1101/2025.09.01.673430

16 Dumoux, M. et al. Cryo-plasma FIB/SEM volume imaging of biological specimens. eLife 12, e83623 (2023). 10.7554/eLife.83623

17 O’Connor, S. et al. Demonstrating soft X-ray tomography in the lab for correlative cryogenic biological imaging using X-rays and light microscopy. Scientific Reports 15, 45491 (2025). 10.1038/s41598-025-29385-5

18 Yuen, E. L. H. et al. Membrane contact sites between chloroplasts and the pathogen interface underpin plant focal immune responses. Plant Cell 37 (2025). 10.1093/plcell/koaf214

19 Downes, K. et al. Multi-scale Molecular Imaging of Human Cells reveals COPI and COPII Vesicles at ER Exit Sites. bioRxiv, 2025.2007.2029.667472 (2025). 10.1101/2025.07.29.667472

20 Mastronarde, D. N. Automated electron microscope tomography using robust prediction of specimen movements. J Struct Biol 152, 36–51 (2005). 10.1016/j.jsb.2005.07.007

21 Grant, T. & Grigorieff, N. Measuring the optimal exposure for single particle cryo-EM using a 2.6 Å reconstruction of rotavirus VP6. eLife 4, e06980 (2015). 10.7554/eLife.06980

22 Zheng, S. et al. AreTomo: An integrated software package for automated marker-free, motion-corrected cryo-electron tomographic alignment and reconstruction. J Struct Biol X 6, 100068 (2022). 10.1016/j.yjsbx.2022.100068

23 Kremer, J. R., Mastronarde, D. N. & McIntosh, J. R. Computer visualization of three-dimensional image data using IMOD. J Struct Biol 116, 71–76 (1996). 10.1006/jsbi.1996.0013

24 Hyun, J. K. et al. Membrane remodeling by the double-barrel scaffolding protein of poxvirus. PLoS Pathog 7, e1002239 (2011). 10.1371/journal.ppat.1002239

25 Condit, R. C., Moussatche, N. & Traktman, P. In a nutshell: structure and assembly of the vaccinia virion. Adv Virus Res 66, 31–124 (2006). 10.1016/s0065-3527(06)66002-8

26 Hashimoto, Y., Sheng, X., Murray-Nerger, L. A. & Cristea, I. M. Temporal dynamics of protein complex formation and dissociation during human cytomegalovirus infection. Nat Commun 11, 806 (2020). 10.1038/s41467-020-14586-5

27 Pražák, V. et al. Molecular plasticity of herpesvirus nuclear egress analysed in situ. Nat Microbiol 9, 1842–1855 (2024). 10.1038/s41564-024-01716-8

28 Close, W. L., Anderson, A. N. & Pellett, P. E. in Human Herpesviruses (eds Yasushi Kawaguchi, Yasuko Mori, & Hiroshi Kimura) 167–207 (Springer Singapore, 2018).

29 Wedemann, L., Flomm, F. J. & Bosse, J. B. The unconventional way out—Egress of HCMV through multiviral bodies. Molecular Microbiology 117, 1317–1323 (2022). 10.1111/mmi.14946

30 Pražák, V., Grünewald, K. & Kaufmann, R. in Methods in Cell Biology 253–271-253–271 (Elsevier, 2021).

31 Falckenhayn, J. et al. On-lamella super-resolution cryo-CLEM for cryo-ET enabled by vacuum-free ultra-stable cryogenic fluorescence microscopy. bioRxiv, 2026.2004.2014.717675 (2026). 10.64898/2026.04.14.717675

32 Ott, F., Jönsson, M. R., Grünewald, K. & Hellert, J. Preparation of Bunyavirus-Infected Cells for Electron Cryo-Tomography. Methods Mol Biol 2824, 221–239 (2024). 10.1007/978-1-0716-3926-9_15

33 Liu, Y. et al. Leonardo: a toolset to correct sample-induced artifacts in light sheet microscopy images. bioRxiv, 2025.2010.2026.684661 (2025). 10.1101/2025.10.26.684661

34 Heumann, J. M., Hoenger, A. & Mastronarde, D. N. Clustering and variance maps for cryo-electron tomography using wedge-masked differences. J Struct Biol 175, 288–299 (2011). 10.1016/j.jsb.2011.05.011

35 Lamm, L. et al. MemBrain v2: an end-to-end tool for the analysis of membranes in cryo-electron tomography. bioRxiv, 2024.2001.2005.574336 (2025). 10.1101/2024.01.05.574336

36 Pettersen, E. F. et al. UCSF ChimeraX: Structure visualization for researchers, educators, and developers. Protein Sci 30, 70–82 (2021). 10.1002/pro.3943

37 Martinez-Sanchez, A., Baumeister, W. & Lučić, V. Statistical spatial analysis for cryo-electron tomography. Computer Methods and Programs in Biomedicine 218, 106693 (2022). 10.1016/j.cmpb.2022.106693

38 Schiøtz, O. H. et al. Serial Lift-Out: sampling the molecular anatomy of whole organisms. Nature Methods 21, 1684–1692 (2024). 10.1038/s41592-023-02113-5

