## Supplementary_information for "View Tomo: Context-aware targeting and analysis in electron cryo-tomography"

### Protocol

The View Tomo script for acquisition in an automated manner is implemented using SerialEM script command interface to enable the collection of low magnification tilt series using so-called View parameters as defined in the "Low Dose Control" menu. The proposed version of the script can run as a single View Tomo acquisition at a given site of interest (see *Single View Tomo Acquisition*) or be used in batch through the "Acquire at Items" dialogue (see *Batch View Tomo Acquisition*). If necessary, refer to the SerialEM website, tutorials and online documentation at https://bio3d.colorado.edu/SerialEM/download.html or Nexperion https://bio3d-mirror.nexperion.net/ftp/SerialEM/.

##### *Importing the Script*

To use the provided version of the View Tomo script, copy the content of the "ViewTomo_batch" file into an empty SerialEM script slot in the script command interface:

1. Open SerialEM
2. Enter the "Script" command interface in the dedicated menu in SerialEM
3. Select "Edit" and choose the number of an empty script slot
4. In parallel, open the "ViewTomo_batch" file with a text editor
5. Then, copy the ViewTomo_batch source code into the script editor and save the script

##### *Preparation*

1. Load cryoEM grid(s) carrying specimen into the transmission electron microscope according to manufacturer's recommended procedure
   *Note: When examining samples prepared by cryo focused ion beam (FIB) milling of thin lamellae in particular, special attention has to be paid to the orientation in which the grids are loaded in the cassette (e.g. for ThermoFisher microscopes) to properly orient the side of the grid, which has the sample on top as well as lamellae perpendicular to the stage tilt axis once loaded inside the column (https://github.com/eisfabian/PACEtomo/blob/main/img/gridOrientation.png)*
2. Acquire grid map at 135-times magnification using the "Setup Full Montage" routine in SerialEM (see https://bio3d.colorado.edu/SerialEM/hlp/html/hidd_montagesetup.htm)
3. Locate regions of interest (ROI) (*e.g.* lamellae) in the grid map and save their position by creating corresponding navigator items using the "Add Points" feature (https://bio3d.colorado.edu/SerialEM/hlp/html/about_navigator.htm)
4. Save the navigator in the drop-down menu
5. Switch to low-dose and use the "Low Dose Control" tab (pink) to configure the low-dose mode called "VIEW" with the desired illumination conditions
   a. *Typically, for FIB-milled cellular specimens, 3600-times was adequate to image lamellae of ~10 µm in length and width.*

b. *Adjust the beam settings to achieve a dose rate of 50 e-px$^{-2}$ measured over vacuum. This results in ~20 e-px$^{-2}$ over most specimens.*
c. *Using the "Set Up" dialogue in the "Camera and Scripts" tab, set the View acquisition parameters to: 1 s exposure time, bin 1, and no dose fractionation.*

#### *Single View Tomo Acquisition*

1. Using the acquired grid map, navigate to a ROI (*e.g.* lamellae)
2. Determine eucentric height by entering the "Tasks" menu and selecting "Rough Eucentricity" (https://bio3d.colorado.edu/SerialEM/hlp/html/menu_tasks.htm).
3. Afterwards, select "Update Z" in the Navigator to save the determined eucentric height for the corresponding navigator item.
   *Note: Additionally running "Fine Eucentricity" is recommended after "Rough Eucentricity" if acquiring View tomograms at 8,700-times magnification or higher.*
   *Note: Automated eucentricity determination can fail for lamella due to the strong contrast at lamella edges or the presence of large ice crystals, in these cases checking and adjusting eucentricity by manually tilting, acquiring an image and adjusting the z-height so that the centre image does not move can be a useful approach.*
4. Enter the "Script" menu in SerialEM. Select "Edit" and then, the script slot where you previously saved the ViewTomo_batch script (see Importing the script).
   *Note: In the ViewTomo_batch source code the following tilt series acquisition parameters defined by the following script variables can be edited to modify the values set by default:*

| **Acquisition parameters** | **Script Variable** | **Default Value** |
|---|---|---|
| Starting tilt angle (in degrees) of the dose-symmetric tilt series. In the case of lamella a pretilt that compensates for the milling angle can be entered (*e.g.* -8) | tiltStart | 0 |
| Maximum tilt angle (in degrees) of the tilt series | tiltRangePos | 51 |
| Minimum tilt angle (in degrees) of the tilt series | tiltRangeNeg | 51 |
| Angle increment step (in degrees) between tilts | tiltStep | 3 |

5. Centre your ROI using the View mode
6. Optional: Open a new file in the "File" menu, select "Open New" and save it in your desired folder.
7. Select "Run" in the script dialogue

#### ***Batch View Tomo Acquisition***

The batch collection procedure relies on the "Acquire at Items" dialogue so each acquisition position needs to be defined in the Navigator beforehand (see Preparation)

1. Using the acquired grid map, navigate to a ROI (*e.g.* lamellae) preselected in the Navigator
2. Determine eucentric height by entering the "Tasks" menu and selecting "Rough Eucentricity" performed directly in SerialEM.
   *Note: Additionally running "Fine Eucentricity" is recommended after "Rough Eucentricity" if acquiring View tomograms at 8,700-times magnification or higher. In case you require "Fine eucentricity" to be run additionally, this can be also implemented in the "Acquire at Items" dialogue.*
3. Update the coordinates for the current ROI using "Move Item" in the navigator so that the exact stage position in X, Y and Z corresponds to the correct coordinates resulting from Rough Eucentricity in the Navigator
4. Define coordinates for the collection of view tomograms by selecting position in the Navigator and ticking "Acquire (A)"
5. Tick "New file at item" this will prompt you to define the base name of the files to be acquired, directory for saving, and open a new file for each item during acquisition
6. Perform steps 1-5 for all desired view tomogram positions
7. Enter the "Navigator" menu in SerialEM and select "Acquire at Items"
8. In the "Primary Action" tab, check "Run script" and select the ViewTomo_batch script in the drop-down menu (see Importing the script)
9. In the "Tasks before or after Primary Action" all check boxes can be unticked although we recommend keeping "Manage Dewars/Vacuum" active.
10. Hit the "Go" button to start
11. Individual tilt series will be saved at each View tomo position and acquired stack of images accessible in the working directory

#### ***Reconstructing tilt series***

1. Export the raw tiltseries data from the microscope to a suitable location for processing. Reconstruction requires a cluster or command line/linux environment.
2. There are 2 main proposed pipelines for View Tomo reconstruction:
   a. Single-step reconstruction:
      i. We provide a script "viewtomo_align" that is a single command line entry that will perform all the steps outlined below resulting in a reconstructed tomogram: https://github.com/vojtaprazak/ViewTomo

      *Note: We particularly recommend this for the on-the-fly processing when aiming for subsequent high magnification tilt series acquisition in a single microscope session*
      *Note: The output from this script is compatible with further processing and refinement of the tomogram alignment and reconstruction in IMOD*
   b. Multi-step reconstruction
      i. Tilts images initially saved as a stack during data collection using the symmetric tilt-scheme, need to be re-sorted from such bidirectional acquisition to follow the order of consecutive tilt angles (-60° to +60° ) prior to reconstruction. Use the newstack function of IMOD to sort the tilt-series

by tilt-angle. To do so, run the command: “newstack -reo 1 inputFile.mrc outputFile.mrc”

ii. For reconstruction of view tomograms, load reordered tilt-series into IMOD Etomo (see https://bio3d.colorado.edu/imod/doc/etomoTutorial.html)

iii. In the “Fiducial Model Generation” tab, select the patch tracking option for tilt-series alignment.
*Note: For FIB-milled specimens, it is recommended to use the “Create Boundary Model” feature to exclude unmilled areas of the sample and vacuum by limiting patch tracking to the lamella.*
*Note: When fiducials are present it is advised to use these for better alignment if the view tomogram is to be used for downstream processing (e.g. segmentation, biological interpretation).*

3. For tomogram generation we recommend implementing ctf-correction of aligned stacks using the estimation from IMOD’s ctfplotter.
*Note: Final tomograms should be reconstructed at bin 2 and trimmed in x,y and z if extending beyond the dimensions of the targeted lamella. This allows for sufficient pixel size for analysis while also saving on data requirements.*
*Note: Optional filtered during reconstruction with a SIRT-like filter equivalent to 15 iterations should be suitable to visualise most cellular features.*
*Note: It is best practice to delete all intermediate files upon completion of tomogram reconstruction to limit the storage requirements per data point.*

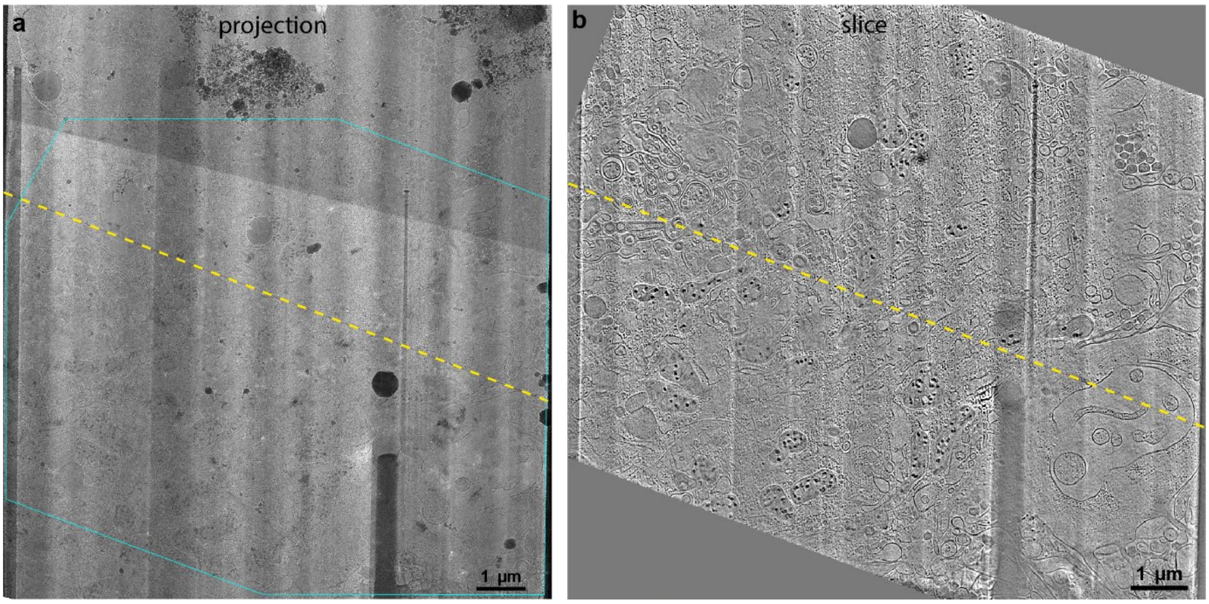


**Supplementary Figure 1. a,** Transmission electron image of a cryo-lamella and **b**, a view tomogram sliceof the same area (panel B). Yellow dashed line represents the tilting axis.  Cyan outline in **a** corresponds to the area in panel B covered by the tomographic acquisition.

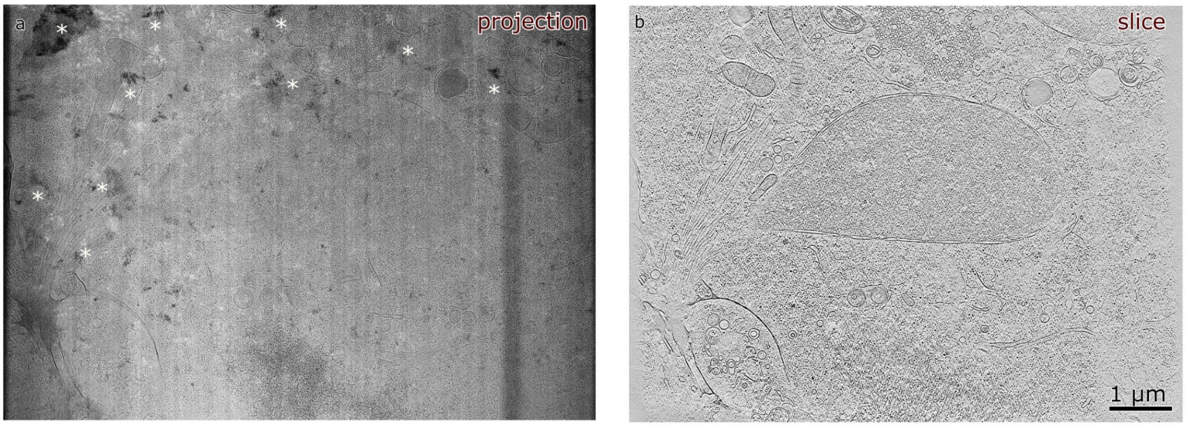


**Supplementary Figure 2. View Tomo data for an RVFV infected cell. a**, Projection image of the region shown in Fig. 5, displaying pronounced Bragg reflections indicative of incomplete vitrification. These features obscure structural detail in the projection image.

**b**, Slice through the corresponding view tomogram used for segmentation in Fig. 5. Despite the presence of crystalline ice, the tomogram remains interpretable and does not exhibit obvious preservation artefacts at this scale.

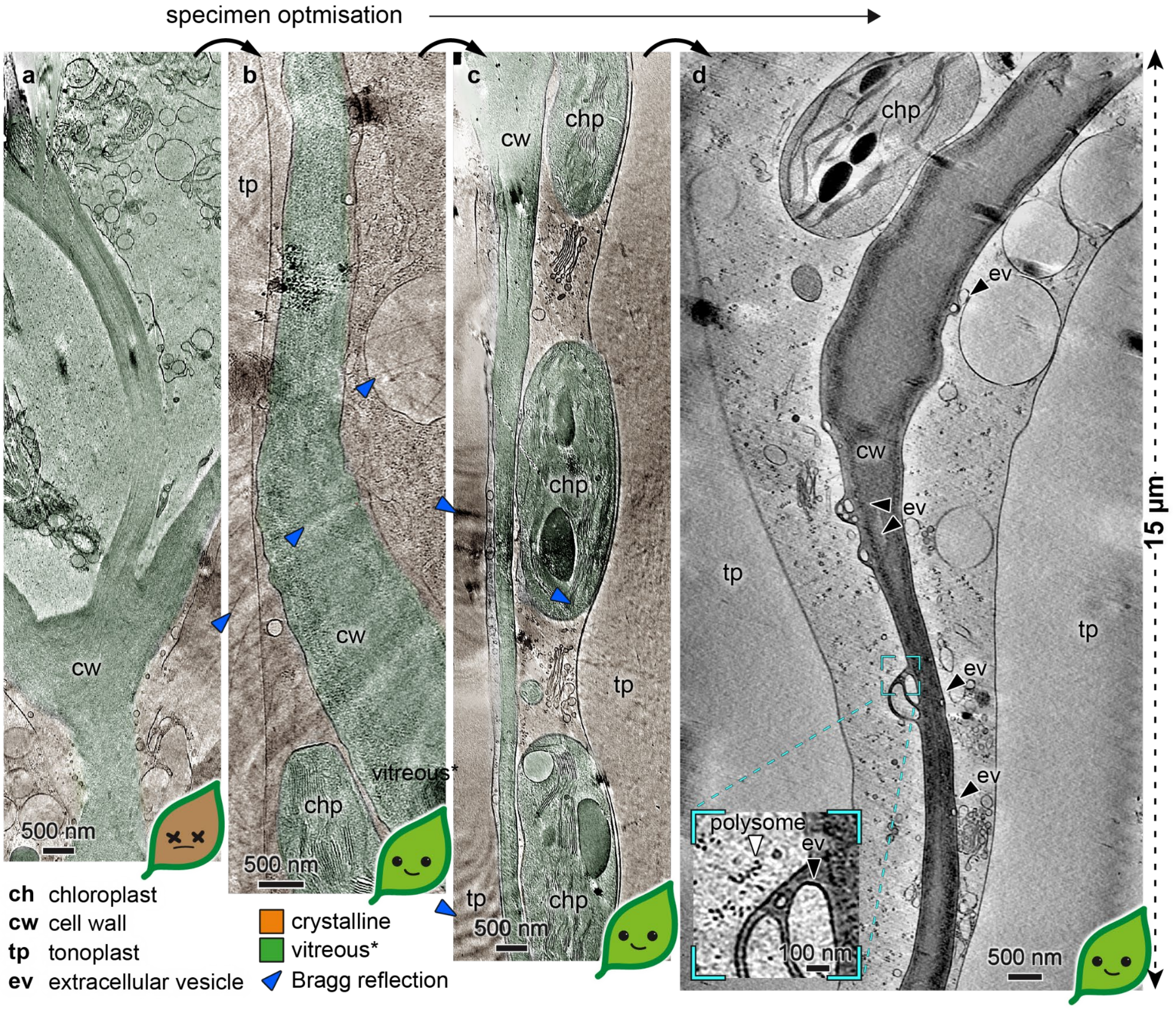


**Supplementary Figure 3. View tomograms enable rapid assessment of biological state and preservation quality during sample preparation and optimisation** Panels **a-d** show sections through view tomograms illustrate a progression in specimen preparation towards **d**, culminating in complete vitrification of a serial-lift out of *Nicotiana benthamiana* leaves. **a**, Epidermal peels were frozen using the waffle method resulting in a wide-spread cell death and partial vitrification which was readily identifiable in view tomograms. **b**, **c**, High-pressure freezing of whole leaf sections using different filler compounds, yielding structurally preserved specimens in which selected compartments (for example, chloroplasts and mitochondria) appeared vitreous, while much of the cytosol and vacuole contains crystalline ice. **d**, Addition of 180 mM glycerol results in fully vitrified specimens suitable for further analysis.

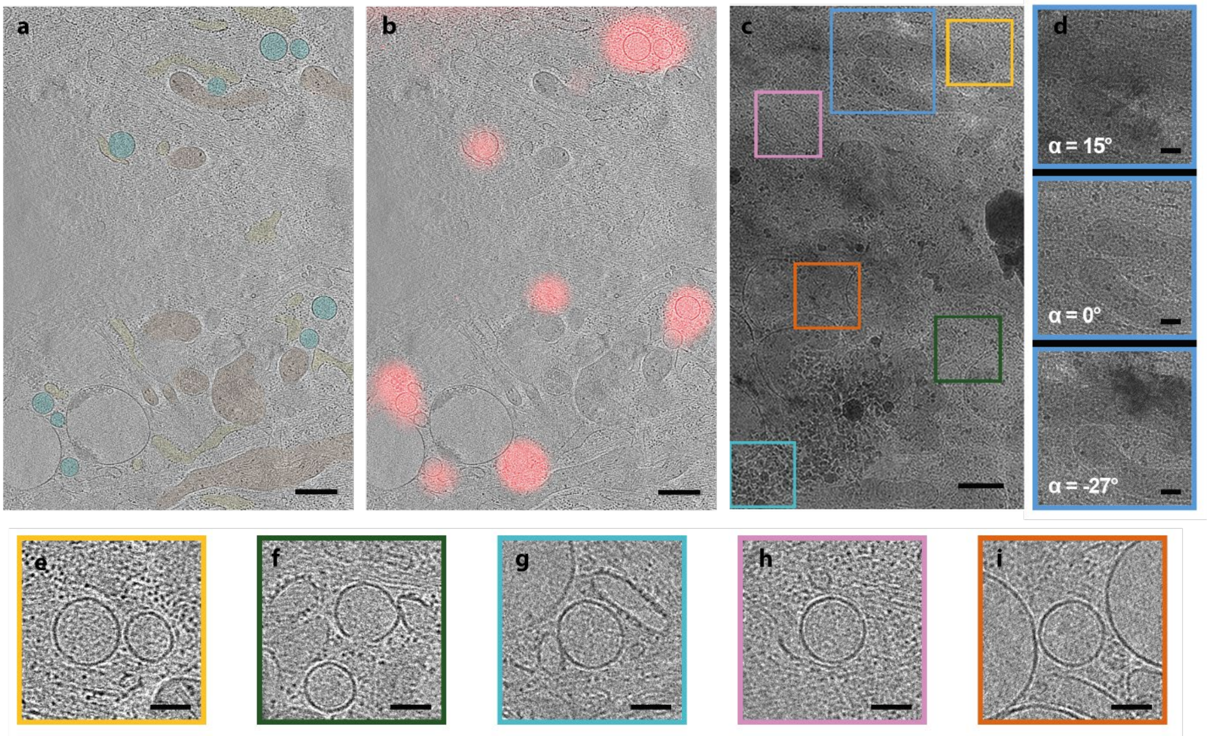


**Supplementary Figure 4. Identification of indistinctive vesicles in overview tomogram of human fibroblasts as peroxisomes using cryoCLEM approach. a,** Slice through an overview tomogram of lamella prepared from a primary MRC5 cell with peroxisomes fluorescently labelled by transient transfection of PTS1-mCherry. Organelles were annotated depicting peroxisomes (turquoise), mitochondria (orange) and endoplasmic reticulum (yellow). Scale bar 500 nm. **b**, CryoCLEM image visualizing superposition of slice through overview tomogram and PTS1-mCherry fluorescence recorded at 550 nm excitation using cryoFM. Image registration was performed using correlation between cryoFM data and a corresponding cryoTEM micrograph of the lamellae recorded at 4,800-times (view) magnification. Scale bar 500 nm. **c**, Projection image of the same lamella at 0° tilt, blue inset indicates the same feature at different stage tilt angles (15, 0 and -27) illustrating the use of View Tomo for gauging sample vitrification by visualising Bragg diffractions, which can be more clearly observed upon tilting the specimen. Scale bar 200 nm. **d**, Magnified sections of different slices through overview tomogram (**b**) from regions depicted in **c** indicate presence and location of peroxisomes based on fluorescence correlation and feature identification in view tomograms. Scale bar 200 nm.

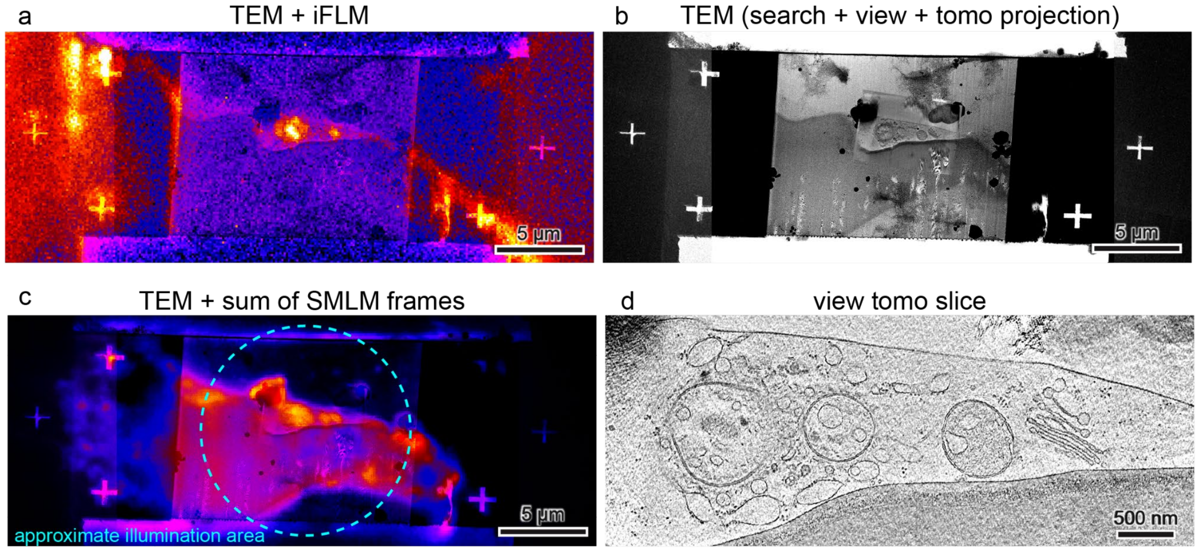


**Supplementary Figure 5. Comparison of cryo-fluorescence imaging before and after View Tomo. a,** Overlay of cryoEM data with cryo-fluorescence microscopy (cryoFM) acquired using the integrated fluorescence microscope (iFLM) within the Aquilos cryoFIB-SEM. Fluorescence data were collected at 10% diode intensity with a 2 s integration time. **b**, CryoEM data only, shown as a composite of a search image, a view image, and a projection through the corresponding view tomogram. Correlation between FM and EM signal was achieved using fiducial markers milled through the lamella. **c**, Overlay of cryoEM data with cryoFM acquired after TEM imaging using a custom cryogenic fluorescence microscope equipped with an ultrastable stage (see Methods). The fluorescence image represents a sum of frames collected over 1.42 hours at an illumination intensity of ~200 W cm$^{-2}$. The approximate illumination area is indicated in cyan. Ice crystals deposited on the surface of the lamella contribute additional fluorescent signal, which accounts for the primary differences between pre- and post-TEM fluorescence images. **d**, Representative slice through the view tomogram corresponding to the region shown in Fig. 6.

#### Supplementary methods:

##### *CryoCLEM-guided HCMV workflow in MRC-5 fibroblasts*

##### *Cell culture and grid preparation*

Primary MRC-5 human fibroblasts were cultured in DMEM containing 10% FBS and 1× GlutaMAX at 37 °C and 5% $CO_2$ until 70–80% confluency. Holey $SiO_2$ film on R 2/2 gold 200 mesh grids were glow discharged for 60 s at negative polarity and 25 mA plasma current. Cell adhesion was facilitated by applying 50 µl fibronectin solution (50 µg $ml^{-1}$ in 20 mM HEPES) to each grid and incubating for 2 h at 37 °C.

For negative-control grids, 6,000 cells were seeded per grid. Grids intended for transfection and/or infection were seeded with 8,000–10,000 cells per grid to compensate for cell loss during downstream handling. Cells were incubated overnight at 37 °C and 5% $CO_2$.

##### *Transfection and infection*

Transient transfection was carried out using Lipofectamine 3000 in serum-free OptiMEM according to the manufacturer's instructions. The transfection mix was adjusted to deliver 5 ng plasmid DNA, 20 µl Lipofectamine 3000 and 10 nl P3000 per 1,000 cells, with a final applied volume not exceeding 50 µl per grid. Cells were incubated for 24 h after transfection.

Cells were infected with HCMV AD169-pUL37x1-GFP at MOI 5 for transfected cells and MOI 3 for untransfected cells. Virus was diluted in serum-free DMEM and applied in a volume not exceeding 50 µl per grid. Infection proceeded for 1 h at 37 °C and 5% $CO_2$.

##### *Vitrification*

Cells on EM grids were plunge-frozen in a liquid ethane/propane mixture at −180 °C using an EM GP2 Automatic Plunge Freezer. Prior to vitrification, grids were washed with serum-free FluoroBrite. Blotting was performed for 7–8 s at 37 °C and 80% humidity.

##### *Gallium cryoFIB milling*

Gallium cryoFIB milling was performed on an Aquilos 2 Cryo-FIB-SEM instrument. After mapping the grids, the specimen was coated sequentially by sputter coating for 10 s at 0.1 mbar and 30 mA, GIS deposition for 70 s, and sputter coating for 5 s at 0.1 mbar and 7 mA. Milling was automated using SerialFIB[1] at a fixed milling angle of 8°. Rectangular fiducial holes were milled adjacent to each suitable lamella at 90°. Rough milling thinned the sample at ROIs to approximately 500 nm. After cryoFM imaging, final polishing to 150–250 nm thickness was performed manually at 10–50 pA using beam shift. A final sputter coat of 5 s at 0.1 mbar and 7 mA was then applied.

##### *Cryo-fluorescence microscopy and image registration*

For gallium cryoFIB-milled samples, cryoFM was performed on a STELLARIS 8 confocal microscope. Prior to cryoFIB milling, widefield mode was used to map fluorescence across the grid and identify candidate lamella sites. Following rough milling, confocal mode was used to visualise fluorescently labelled features within lamellae by recording sequential images at 50× magnification over 5–6 µm in z with 300 nm spacing.

Tile sets of view images were collected for each viable lamella, stitched in Adobe Photoshop, and registered to the corresponding cryoFM data using Icy ec-CLEM[2,3]. These correlations were used to identify regions for high-resolution tomography.

##### *View Tomo and high-resolution cryoET*

View tomograms were acquired at 3,600-times to 4,800-times magnification using the ViewTomo_batch script in batch mode. Low-magnification tilt series were collected bi-directionally from 0° to ±51° using 3° increments and 2 s exposure at bin 1.

Once view tomograms had been reconstructed and correlated with cryoFM data, high-magnification tilt series were set up to target regions of interest identified by cryoCLEM. High-magnification tilt series were typically acquired from +68° to −52° with 3° increments, a target defocus of −5 to −7 µm, and a total electron dose of approximately 140–180 $e^{-}$ $Å^{-2}$ per tilt series.

##### *Reconstruction*

High-magnification tilt series were aligned by marker-free alignment in AreTomo and reconstructed in IMOD/Etomo. Low-magnification view tomograms were reconstructed in IMOD/Etomo using patch tracking.

**Supplementary text 1, ViewTomo_batch SerialEM script**

This script can also be found online in the SerialEM script repository: https://serialemscripts.nexperion.net/script/88

```
# Name:        ViewTomo_batch
# Author:    Robert Gebauer
# Version:    1.0
# Last update:  19.03.2026
# Support:   

### Tilt series parameters

tiltStart    = 0
tiltRangePos    = 51
tiltRangeNeg    = 51
tiltStep    = 3

### Default settings

tiltLimit    = 68
tiltBacklash    = -1
wait        = 3

### Initialization

tiltNum = 0
currentTiltAngle = 0
ReportStageXYZ
stageX = $ReportedValue1
stageY = $ReportedValue2
stageZ = $ReportedValue3
currentTiltAngle = $tiltStart

GoToLowDoseArea V

### Image acquisition at tiltStart

TiltTo $tiltStart
View
Save

### Image acquisition for positive tilt angles

tiltNum = $tiltRangePos / $tiltStep
```

```
loop $tiltNum indexP
  currentTiltAngle = $tiltStart + ( $tiltStep * $indexP )
  if $currentTiltAngle > ( $tiltStart + $tiltRangePos )
    break
  elseif $currentTiltAngle <= $tiltStart
    break
  elseif $currentTiltAngle > $tiltLimit
    break
  endif
  tiltTo $currentTiltAngle
  MoveStageTo $stageX $stageY $stageZ
  delay $wait sec
  View
  Save
endloop

### Image acquisition for negative tilt angles

currentTiltAngle = 0
tiltNum = $tiltRangeNeg / $tiltStep
loop $tiltNum indexR
  currentTiltAngle = $tiltStart - ( $tiltStep * $indexR )
  if $currentTiltAngle < ( $tiltStart - $tiltRangeNeg )
    break
  elseif ( $currentTiltAngle + $tiltLimit ) < 0
    break
  elseif $currentTiltAngle >= $tiltStart
    break
  endif
  tiltTo $currentTiltAngle
  tiltBy $tiltBacklash
  tiltTo $currentTiltAngle
  MoveStageTo $stageX $stageY $stageZ
  delay $wait sec
  View
  Save
endloop

### Finishing up

CloseFile
TiltTo 0
```

#### Supplementary References


1 Klumpe, S., Fung, H. K. H., Goetz, S. K., Zagoriy, I., Hampoelz, B., Zhang, X., Erdmann, P. S., Baumbach, J., Müller, C. W., Beck, M., Plitzko, J. M. & Mahamid, J. A modular platform for automated cryo-FIB workflows. *eLife* **10**, e70506 (2021). https://doi.org/10.7554/eLife.70506

2 de Chaumont, F., Dallongeville, S., Chenouard, N., Hervé, N., Pop, S., Provoost, T., Meas-Yedid, V., Pankajakshan, P., Lecomte, T., Le Montagner, Y., Lagache, T., Dufour, A. & Olivo-Marin, J.-C. Icy: an open bioimage informatics platform for extended reproducible research. *Nature Methods* **9**, 690-696 (2012). https://doi.org/10.1038/nmeth.2075

3 Paul-Gilloteaux, P., Heiligenstein, X., Belle, M., Domart, M.-C., Larijani, B., Collinson, L., Raposo, G. & Salamero, J. eC-CLEM: flexible multidimensional registration software for correlative microscopies. *Nature Methods* **14**, 102-103 (2017). https://doi.org/10.1038/nmeth.4170
